# RNA molecules display distinctive organization at nuclear speckles

**DOI:** 10.1101/2022.10.17.512423

**Authors:** Sneha Paul, Mauricio A. Arias, Li Wen, Susan E. Liao, Jiacheng Zhang, Xiaoshu Wang, Oded Regev, Jingyi Fei

**Affiliations:** Department of Biochemistry and Molecular Biology, The University of Chicago, Chicago, IL, 60637; Courant Institute of Mathematical Sciences, New York University, New York, NY, 10012; Department of Physics, The University of Chicago, Chicago, IL, 60637; Graduate Program in Biophysical Sciences, The University of Chicago, Chicago, IL, 60637; The College, The University of Chicago, Chicago, IL, 60637; Institute for Biophysical Dynamics, The University of Chicago, Chicago, IL, 60637

## Abstract

RNA molecules often play critical roles in assisting the formation of membraneless organelles in eukaryotic cells. Yet, little is known about the organization of RNAs within membraneless organelles. Here, using super-resolution imaging and nuclear speckles as a model system, we demonstrate that different sequence domains of RNA transcripts exhibit differential spatial distributions within speckles. Specifically, we image transcripts containing a region enriched in SR protein binding motifs and another region enriched in hnRNP binding motifs. We show that these transcripts localize to the outer shell of speckles, with the SR motif-rich region localized closer to the speckle center relative to the hnRNP motif-rich region. Further, we identify that this intra-speckle RNA organization is driven by the strength of RNA-protein interactions inside and outside speckles. Our results hint at novel functional roles of nuclear speckles and likely other membraneless organelles in organizing RNA substrates for biochemical reactions.

## Introduction

Eukaryotic cells contain many membraneless organelles with distinct nuclear (*1*, *2*) or cytoplasmic localizations (*3*, *4*). These membraneless organelles generally contain RNAs, RNA binding proteins (RBPs) and ribonucleoprotein (RNP) assemblies (*5*, *6*). Multivalent interactions between protein and RNA components drive the formation of these organelles through phase separation (*7–11*), and can also lead to the formation of sub-domains or layered structures within many of them (*12*).

Little is known about the organization of RNA within membraneless organelles, despite its potential role in coordinating biochemical reactions. One exception is the 22 kb long noncoding RNA (lncRNA) *NEAT1*, which serves as a scaffold component of paraspeckles, and is organized with its 5’- and 3’-ends at the paraspeckle shell and its central region at the paraspeckle core (*13*). However, most RNA is not considered a scaffold component. For such non-scaffold, or client RNA transcripts, different localizations of transcripts around or within some membraneless organelles (such as germ granules, nuclear speckles, and stress granules) have been noted (*14–16*). These studies, however, report the localization of RNA transcripts as one entity; whether there is any organization at the level of individual molecules, i.e., between different sequence domains of the same client RNAs, is not clear. Another shortcoming of most previous work is the lack of mechanistic insight explaining the observed localization.

We reasoned that the differential proteome composition inside and outside of a membraneless organelle can lead to a distinctive organization of RNA molecules. Specifically, regions of RNA transcripts interacting with proteins inside the organelle will tend to localize closer to the center than regions interacting with proteins outside the organelle. In this way, the position of RNA transcripts will be driven to the outer shell of the membraneless organelle, and their orientation will be constrained **(Figure 1)**. Here, we refer to the position and orientation of RNA transcripts collectively as intra-organelle RNA organization.

**Figure 1.**
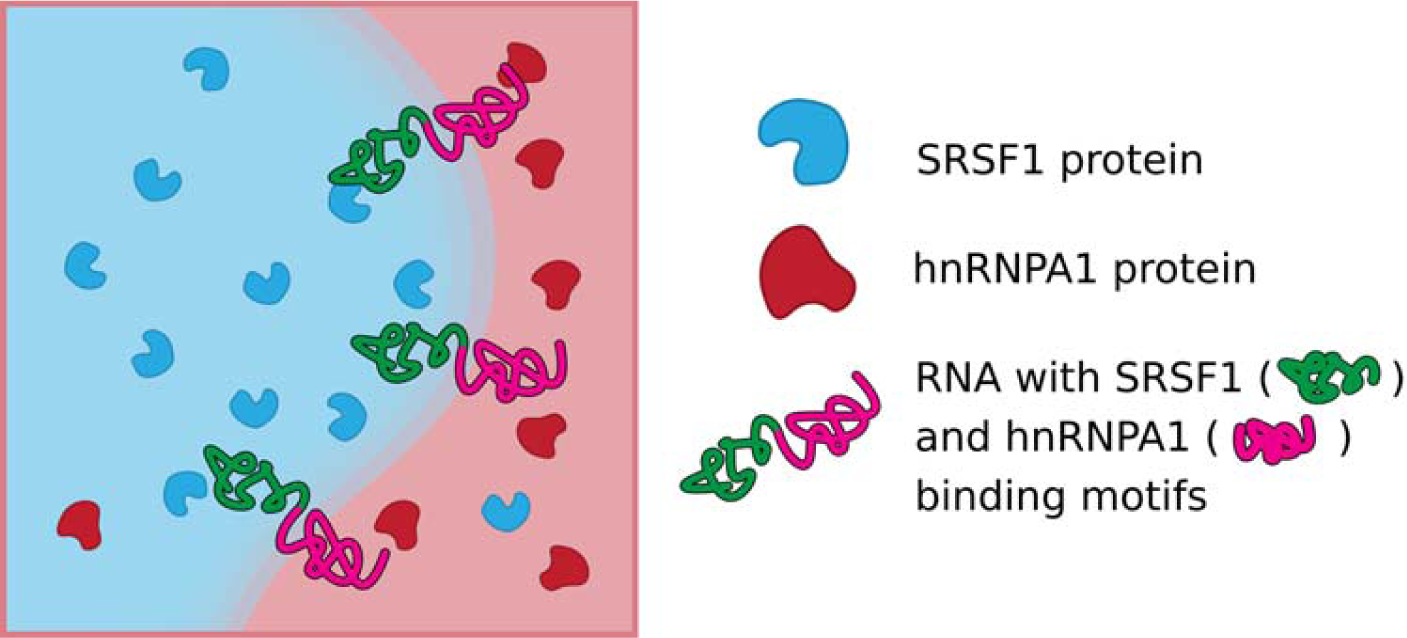
Intra-organelle RNA organization model with nuclear speckle as an example. Nuclear proteins show differential localization with respect to nuclear speckles, with SRSF1 protein enriched in speckles and hnRNPA1 protein distributed in the nucleoplasm. As a result, RNAs containing both RBP binding motifs will be driven to the outer shell of nuclear speckles and will be organized with their SRSF1-binding motifs closer to the speckle center than their hnRNPA1-binding motifs.

We sought to test this intra-organelle RNA organization model using nuclear speckles as a model system. Nuclear speckles are a type of membraneless organelle in higher eukaryotic cells, playing important roles in regulating transcription, splicing and RNA processing (*17*–*20*), and maintaining 3D-genome organization (*21*). Their number ranges between 20 and 50 per cell (*22*) and their size varies from a few hundred nanometers to a few microns (*22*). Nuclear speckles are rich in small nuclear RNAs (snRNAs), spliceosomal proteins, and certain splicing factors, including SR proteins (a family of RBPs named for containing regions with repetitive serine and arginine residues). Polyadenylated RNAs, stained with fluorescently labeled polyT oligos, are observed to localize to nuclear speckle (*23*, *24*). Moreover, a recent transcriptomic analysis has systematically identified nuclear speckle-localized transcripts (*25*). Nuclear speckles exhibit a core-shell organization. Specifically, the scaffold proteins SON and SRRM2 form the core layer of speckles, while spliceosomal components, including snRNAs and spliceosomal proteins as well as the nuclear speckle-localized lncRNA *MALAT1,* are enriched in the outer shell (*16*). In contrast to paraspeckles, nuclear speckles are not known to depend on any specific RNA transcript for their formation.

Nuclear speckles are well suited as a model system for two reasons. First, proteins with different localization relative to nuclear speckles were previously noted. Specifically, certain SR proteins are enriched in nuclear speckles (*26*, *27*), whereas certain heterogeneous nuclear ribonucleoprotein (hnRNP) do not exhibit any enrichment or might be depleted from speckles (*28*, *29*). Second, synthetic reporter constructs can be designed to generate nuclear speckle-localized RNA transcripts (*30*). In this work, we use super-resolution microscopy to demonstrate that the intra-organelle RNA organization model applies to nuclear speckles **(Figure 1)**.

## Results

### SRSF1 and hnRNPA1 proteins exhibit distinct localization relative to nuclear speckles

Confirming previous results, we found that SRSF1 protein was consistently enriched in nuclear speckles (*25*, *26*, *31*) **(Figure S1a)**, whereas hnRNPA1 proteins showed a lower abundance in most nuclear speckles than the surrounding nucleoplasm (*32*) **(Figure S1b)**. In summary, SRSF1 and hnRNPA1 concentrations inside nuclear speckles are distinct from those outside **(Figure S1c)**, justifying the choice of this SR-hnRNP protein pair for further analysis.

### RNAs containing SRSF1- and hnRNPA1-binding motifs display specific intra-speckle organization

We started by synthesizing a reporter construct (MUT_S1-H1_) using the Tet-responsive promoter-controlled three-exon design based on the Chinese hamster *DHFR* gene from our earlier work (*33*). Previous studies indicate that exonic regions are more frequently enriched in SRSF1 binding motifs while intronic regions are more enriched in hnRNPA1 binding motifs (*34*, *35*). Therefore, we included multiple SRSF1-binding motifs in the middle exon and multiple hnRNPA1-binding motifs in the downstream intron **(Figure 2a)**. We verified using BLAST that these sequences exhibit minimal homology with endogenous sequences. Moreover, to ensure that we were imaging RNA transcripts containing both SRSF1 and hnRNPA1 motifs, rather than spliced RNA products, we introduced a mutation at the 3’ splice site of the second intron (AG>GG) which inhibits its splicing (*36*, *37*). The construct was transfected into a HeLa cell line with stably expressed Tet-regulated transactivator Tet-On 3G, and RNA expression was induced with doxycycline. Using a Reverse Transcription-Polymerase Chain Reaction (RT-PCR) assay, we verified that the second intron of MUT_S1-H1_ RNA was mostly unspliced **(Figure S2a, S3a)**, leaving the SRSF1- and hnRNPA1-binding motifs on the same RNA.

**Figure 2.**
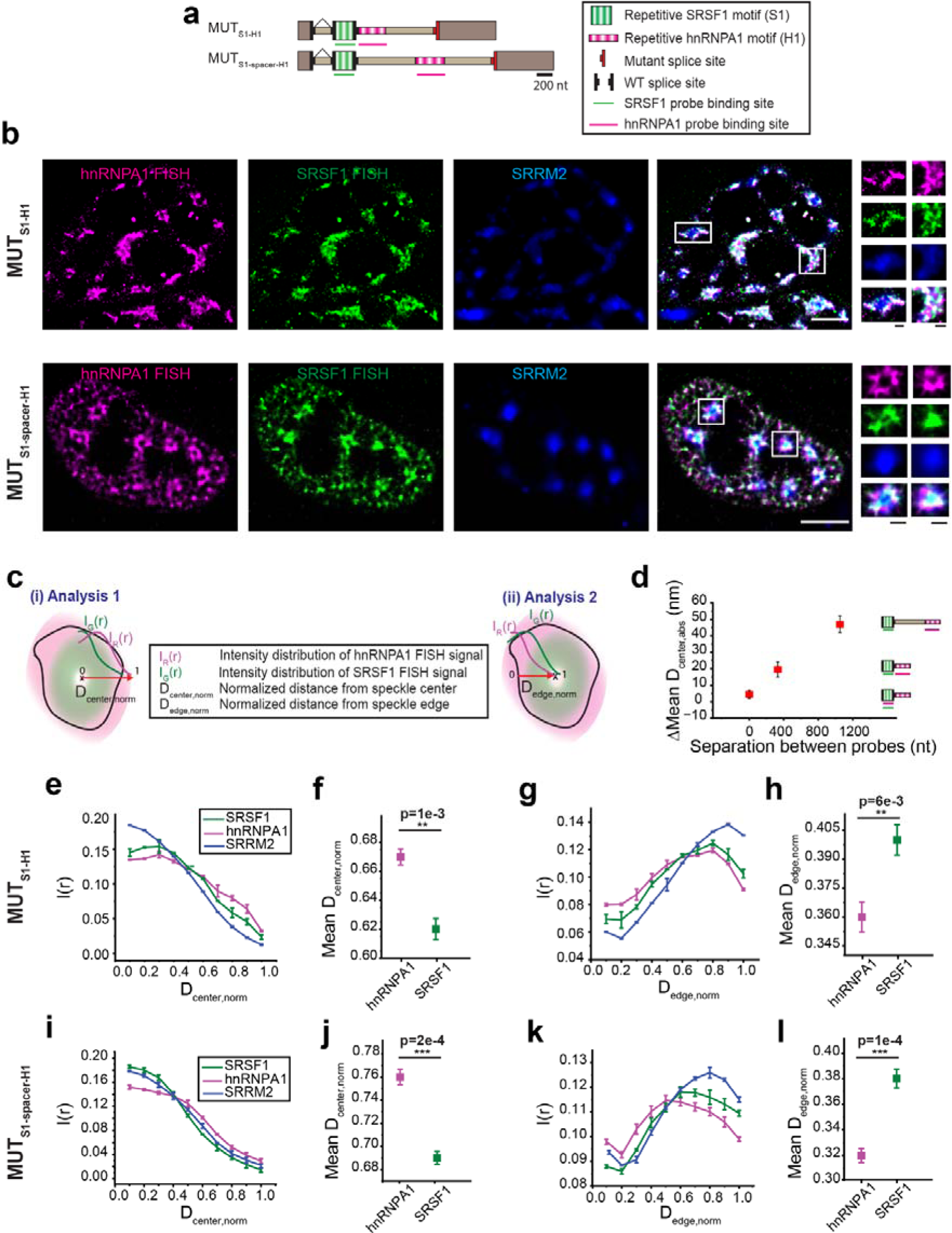
SMLM imaging and analysis of intra-speckle organization of RNAs containing SRSF1 motifs in exon and hnRNPA1 motifs in intron. (a) Schematic illustration of MUT_S1-H1_ and MUT_S1-spacer-H1_ constructs. (b) Representative SMLM image of MUT_S1-H1_ and MUT_S1-spacer-H1_. FISH signals corresponding to hnRNPA1 (labeled with AF647) and SRSF1 (labeled with CF568) motifs in the RNAs are shown in magenta and green respectively. Immunostaining of SRRM2 is shown in blue. Scale bar represents 5 μm (white) and 1 μm (black). (c) Calculation of the distribution of FISH signal as a function of the distance from the center of the nuclear speckle (i) and edge of the nuclear speckle (ii). Due to size differences among nuclear speckles, distances are all normalized from the center of the speckle (i) or the edge of the speckle (ii) to build the overlaid distribution. (d) Plot of difference in absolute mean distance vs. separation in the RNA length (in the unit of nucleotide) between the probe targeting positions. Population distribution of SRSF1 and hnRNPA1 motif signals for MUT_S1-H1_ as a function of the normalized distance from the center of the speckle (e) and edge of the speckle (g). Population-weighted mean normalized distance of SRSF1 and hnRNPA1 signal from the center of speckle (f) and edge of speckle (h) for each speckle for MUT_S1-H1_. Population distribution of SRSF1 and hnRNPA1 motif signals for MUT_S1-spacer-H1_ as a function of the normalized distance from the center of the speckle (i) and edge of the speckle (k). Population-weighted mean normalized distance of SRSF1 and hnRNPA1 signal from the center of speckle (j) and edge of speckle (l) for each speckle for MUT_S1-spacer-H1._ Error bars in the population vs. distance plots report the standard deviation from two replicates, each replicate containing at least 60-90 nuclear speckles collected from 4-6 cells. Scatter plots are generated by combining all nuclear speckles (120–180) from two replicates. Values in scatter plot represent mean ± standard error of mean (s.e.m). p-values in the scatter plots are calculated with paired sample Wilcoxon signed rank test (one-sided), with *p<5e-2, **p<1e-2, ***p<1e-3.

Using fluorescence in situ hybridization (FISH), we labeled the SRSF1 and hnRNPA1-binding motifs on the RNA transcripts with CF568 and Alexa Fluor 647 (AF647), respectively. Nuclear speckles were stained with Alexa Fluor 488 (AF488) labeled antibody against the scaffold protein SRRM2 (*38*). Epifluorescence imaging confirmed that MUT_S1-H1_ RNAs are localized in nuclear speckles **(Figure S3a)**. In addition, minimal non-specific binding of the RNA FISH probes was detected using cells with unsuccessful transfection as controls **(Figure S3b)**.

We next sought to determine the organization of MUT_S1-H1_ RNA in nuclear speckles. One challenge is the small distance expected between the SR and hnRNP motifs. Indeed, since both are on the same RNA transcript and separated by only ∼400 nt, we expect a distance not greater than 50 nm (e.g., the typical physical size of a regular-sized RNA molecule (hundreds to a few thousand nucleotides) is ∼40-100nm, as characterized in solution and in fixed cells (*39*, *40*)). We therefore performed super-resolution imaging on MUT_S1-H1_ RNA using single molecule localization microscopy (SMLM), with a spatial resolution of 10-30 nm (*41*). An epifluorescence image of SRRM2 was recorded before the SMLM imaging on the same cell.

As expected, even with the increased resolution, the resulting images showed rather subtle differences between the two FISH signals at the level of individual speckles **(Figure 2b)**. To obtain statistically significant results, we therefore developed a data analysis pipeline **(Figure S4)** that averages the results across a population of nuclear speckles (∼50-90 per replicate). Specifically, we first selected nuclear speckles containing associated RNA signals by using intensity thresholding on the sum of all three channels, namely the two RNA FISH signal channels and the nuclear speckle marker channel **(Figure S4,** (**1**)). Extremely small, large and irregular nuclear speckles were excluded from the analysis through size and ellipticity cutoffs **(Figure S4,** (**2**)). The radial signal intensity distributions from the two RBP-binding motifs in the RNA were then determined as a function of the normalized distance from the geometric center of the speckle (D_center,norm_, **Figure 2c, (i)**) for individual **speckles (Figure S4,** (**3-4**)). The normalized radial distributions were then averaged across the population of speckles **(Figure S4,** (**5**)). Strikingly, this analysis demonstrated in a statistically significant way that the SRSF1 motif-rich region is distributed closer to the center of the speckle compared to the hnRNPA1 motif-rich region **(Figure 2e-f)**.

In an alternative analysis, we calculated the radial intensity distributions of the two RNA signals as a function of the normalized distance from the edge of the speckle (D_edge,norm_, **Figure 2c, (ii)**, **Figure S4,** (**4-5**)). In agreement with our first analysis, this analysis demonstrated that the SRSF1 motif-rich region is distributed further away from the edge of the speckle, i.e., closer to the center of the speckle, compared to the hnRNPA1 motif-rich region **(Figure 2g-h)**.

Finally, we also estimated the mean absolute radial distance difference between the two regions (D_center,abs_ and D_edge,abs_). We found this distance to be 15-25 nm (Figures 2d and S5), consistent with the typical physical size of an RNA molecule (∼40-100 nm) (*39*, *40*).

Together, these analyses suggest that different sequence domains in an RNA molecule can exhibit differential spatial distributions in nuclear speckles, in agreement with the intra-speckle organization model **(Figure 1)**.

### Validation of intra-speckle RNA organization

We performed several experiments to validate the intra-speckle RNA organization. First, to rule out imaging artifacts due to the choice of the fluorophores, we reversed the FISH labelling scheme, by labeling SRSF1 motifs with AF647 and hnRNPA1 motifs with CF568. We observed the same intramolecular organization trend, using both analysis methods **(Figure S6a-d)**.

Next, we tested whether the measured mean radial distance difference between the two regions can quantitatively reflect the physical distance between the two labeled regions on the transcript, given our imaging resolution. We quantified the radial distribution and absolute mean radial distance for two additional cases: (1) We mixed probes labeled with AF647 and CF568 to target SRSF1 motifs. In this case, we did not observe any significant difference between the radial distribution of AF647 and CF568 signals targeting the same region on the transcript, yielding a near zero (4.5±2.5 nm) difference in the mean radial distance difference between the signals (**Figures 2d, and S5**). (2) We designed MUT_S1-spacer-H1_ by introducing a neutral 720-nt spacer between the SRSF1 and hnRNPA1 motif-rich regions (**Figure 2a**). Introduction of the spacer did not change the splicing outcome **(Figure S2b)**. MUT_S1-spacer-H1_ demonstrated the same intra-speckle organization as MUT_S1-H1_ **(Figure 2i-l)**, with a higher mean radial distance difference of 30-50 nm between the SRSF1 motif-rich and hnRNPA1 motif-rich regions (**Figures 2d and S5**). In summary, the absolute mean radial distance difference between the two labeled regions increased as the number of nucleotides between the two labeled regions increased (**Figure 2d**), confirming that our imaging resolution allows quantification of the physical distance between the two labeled regions on the transcript.

Finally, MUT_S1-H1_ contains repeat sequences of SR and hnRNP binding motifs. To rule out the possibility that the intra-speckle RNA organization is an artifact of these repetitions, we synthesized construct MUT_S1-H1,NR_ **(Figure S7a),** where we randomly mutated 30% of the nucleotides in the SR and hnRNP motif-rich regions, resulting in ‘non-repeat’ (NR) sequences. This construct is still enriched in SRSF1 motifs in the exon and in hnRNPA1 motifs in the intron **(Figure S7b-c)**, but devoid of repeats, as measured by BLAST. RNA transcripts from this construct demonstrated the same splicing behavior as MUT_S1-H1_ **(Figure S2a)**. Importantly, they also demonstrated similar intra-speckle RNA organization **(Figure S7d-g)**. These results confirmed that the intra-speckle RNA organization is driven by the presence of SRSF1 and hnRNPA1 binding motifs, and not because of the sequence repetition.

Together, these results further validate that RNAs exhibit distinctive intra-speckle organization, with SRSF1 motif-rich regions positioned closer to the speckle center compared with hnRNPA1 motif-rich regions.

### Unspliced pre-mRNA exhibits similar intra-speckle organization

We next examined whether transcripts containing a complete set of splice sites, and are thus more similar to endogenous pre-mRNA transcripts, exhibit similar intra-speckle organization. Specially, we replaced the mutated 3’ splice site with an active 3’ splice site to generate the WT_S1-H1_ construct **(Figure 3a)**. WT_S1-H1_ underwent normal splicing and export **(Figure S2c)**. At 30 min induction, the FISH signals for both SRSF1 and hnRNPA1 motifs from the WT_S1-H1_ construct were mostly localized to nuclear speckles (**Figure S3c**), as observed under epifluorescence imaging. At 2 h induction, a significant portion of the FISH signal for the SRSF1 motifs was localized to the cytoplasm, corresponding to the spliced and exported mRNAs, whereas the residual nuclear-localized signals from the SRSF1 and hnRNPA1 motifs remained localized to nuclear speckles, likely corresponding to the pre-mRNA transcripts (**Figure S3c**). This result illustrates that unspliced pre-mRNA can also localize to nuclear speckles similar to the partially spliced RNA.

**Figure 3.**
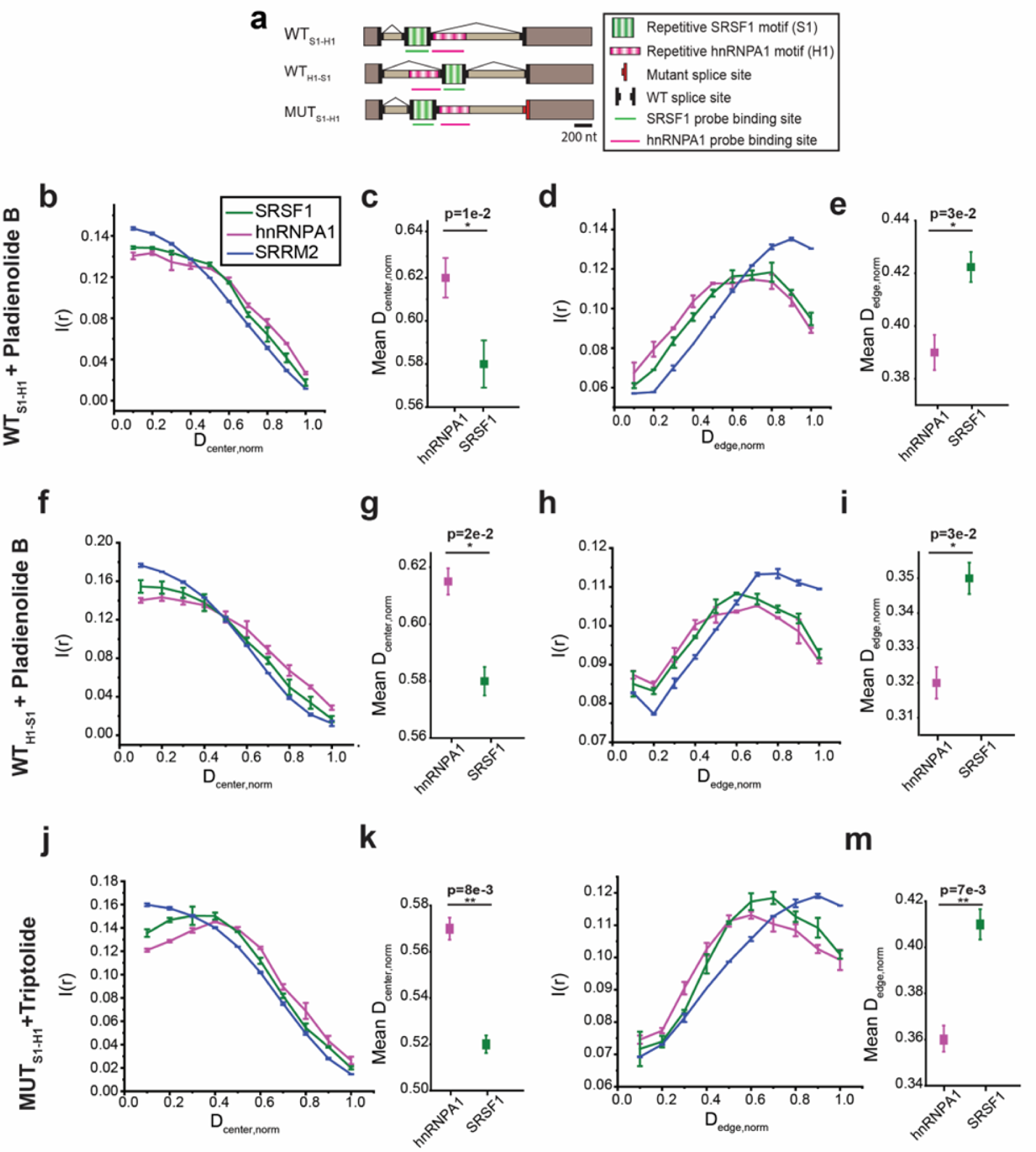
Effect of restored 3’ active splice site and transcription on intra-speckle organization of RNAs containing SRSF1 motifs in exon and hnRNPA1 motifs in intron. (a) Schematic illustration of WT_S1-H1_, WT_H1-S1_ and MUT_S1-H1_. Population distribution of SRSF1 and hnRNPA1 motif signals as a function of the normalized distance from the center of the speckle (b) and edge of the speckle (d) for WT_S1-H1_ in the presence of Pladienolide B. Population-weighted mean normalized distance of SRSF1 and hnRNPA1 signal from the center of speckle (c) and edge of the speckle (e) for each speckle for WT_S1-H1_ in the presence of Pladienolide B. Population distribution of SRSF1 and hnRNPA1 motif signals as a function of the normalized distance from the center of the speckle (f) and edge of the speckle (h) for WT_H1-S1_ in the presence of Pladienolide B. Population-weighted mean normalized distance of SRSF1 and hnRNPA1 signal from the center of speckle (g) and edge of the speckle (i) for each speckle for WT_H1-S1_ in the presence of Pladienolide B. Population distribution of SRSF1 and hnRNPA1 motif signals as a function of the normalized distance from the center of the speckle (j) and edge of the speckle (l) for MUT_S1-H1_ in the presence of Triptolide. Population-weighted mean normalized distance of SRSF1 and hnRNPA1 signal from the center of speckle (k) and edge of the speckle (m) for each speckle MUT_S1-H1_ in the presence of Triptolide. Error bars in the population vs. distance plots report the standard deviation from two replicates, each replicate containing at least 36-48 nuclear speckles collected from 3-4 cells. Scatter plots are generated by combining all nuclear speckles (72–96) from two replicates. Values in scatter plot represent mean ± standard error of mean (s.e.m). p-values in the scatter plots are calculated with paired sample Wilcoxon signed rank test (one-sided), with *p<5e-2, **p<1e-2, ***p<1e-3.

The presence of spliced intron lariat or mature mRNA in the nuclear speckle-localized transcripts can complicate interpretation of the imaging results. As we cannot completely rule out their presence, we inhibited splicing with Pladieonolide B **(Figure S2c, S3c)** (*42*). Splicing inhibition ensures that we image pre-mRNAs containing both SRSF1 and hnRNPA1 motifs. Since splicing inhibition can lead to an enlargement of nuclear speckles (*42*), we compared the morphology of nuclear speckles in the absence and presence of Pladieonolide B **(Figure S8)**. While speckle size increased in the presence of Pladieonolide B, the ellipticity and regularity of the speckle surface were not significantly affected **(Figure S8)**. We applied the same speckle size and ellipticity cutoffs for an unbiased comparison of samples. We found that the SRSF1 motif and hnRNPA1 motif-rich regions of WT_S1-H1_ pre-mRNA exhibit the same intra-speckle organization as that of MUT_S1-H1_ RNA **(Figure 3b-e).**

### Intra-speckle organization is not due to transcription order

Actively transcribed genes are found to be associated with speckle periphery (*17*). One possible explanation for the organization we observed is that the hnRNPA1 motif-rich region, being downstream of the SRSF1 motif-rich region, is transcribed later and is therefore closer to the transcription site positioned outside nuclear speckles. However, we expect that at 2 h after induction, most transcripts are no longer associated with the DNA, and that any orientation that is present during transcription cannot persist for that long.

Nevertheless, to rule out transcription order as a cause, we performed two additional experiments. In the first experiment, we designed construct WT_H1-S1_ in which the hnRNPA1-binding motifs were moved to the intron upstream of the SRSF1 motif-containing middle exon (WT_H1-S1_, **Figure 3a)**. The WT_H1-S1_ construct was spliced similarly to the WT_S1-H1_ construct **(Figure S2d),** and we imaged the pre-mRNA from this construct in the presence of Pladienolide B. In the second experiment, we imaged MUT_S1-H1_ RNA in the presence of the transcription inhibitor, Triptolide. The speckles became more rounded in the presence of Triptolide **(Figure S8)**, as expected upon transcription inhibition (*22*). We again applied the same speckle size and ellipticity cutoffs for an unbiased comparison of samples.

Importantly, in both cases we observed the same intra-speckle RNA organization, in which the SRSF1 motif-rich region is closer to the speckle center than the hnRNPA1 motif-rich region **(Figure 3f-m)**. These results provide strong evidence that the intra-speckle organization is independent of the order in which the SRSF1 motif-rich and hnRNPA1 motif-rich regions are transcribed.

### RNA transcripts with a combination of SRSF7 and hnRNPA1 motifs exhibit similar intra-speckle organization

To test whether RNA containing other combinations of SR and hnRNP motifs exhibits similar positioning and orientation within nuclear speckles, we replaced the SRSF1 motifs in MUT_S1-H1_ and WT_S1-H1_ with SRSF7 motifs to generate MUT_S7-H1_ and WT_S7-H1_ **(Figure 4a)**. The splicing behavior of these constructs was similar to the SRSF1 motif-containing ones **(Figure S2a,c)**. The radial intensity distributions with respect to the center and edge showed that SRSF7 motif-rich region is closer to the center of nuclear speckles than the hnRNPA1 motif-rich region **(Figure 4b-i),** the same trend observed for MUT_S1-H1_ and WT_S1-H1_. These results demonstrate that the intra-speckle organization of SR and hnRNP motifs is not specific to the SRSF1-hnRNPA1 combination.

**Figure 4.**
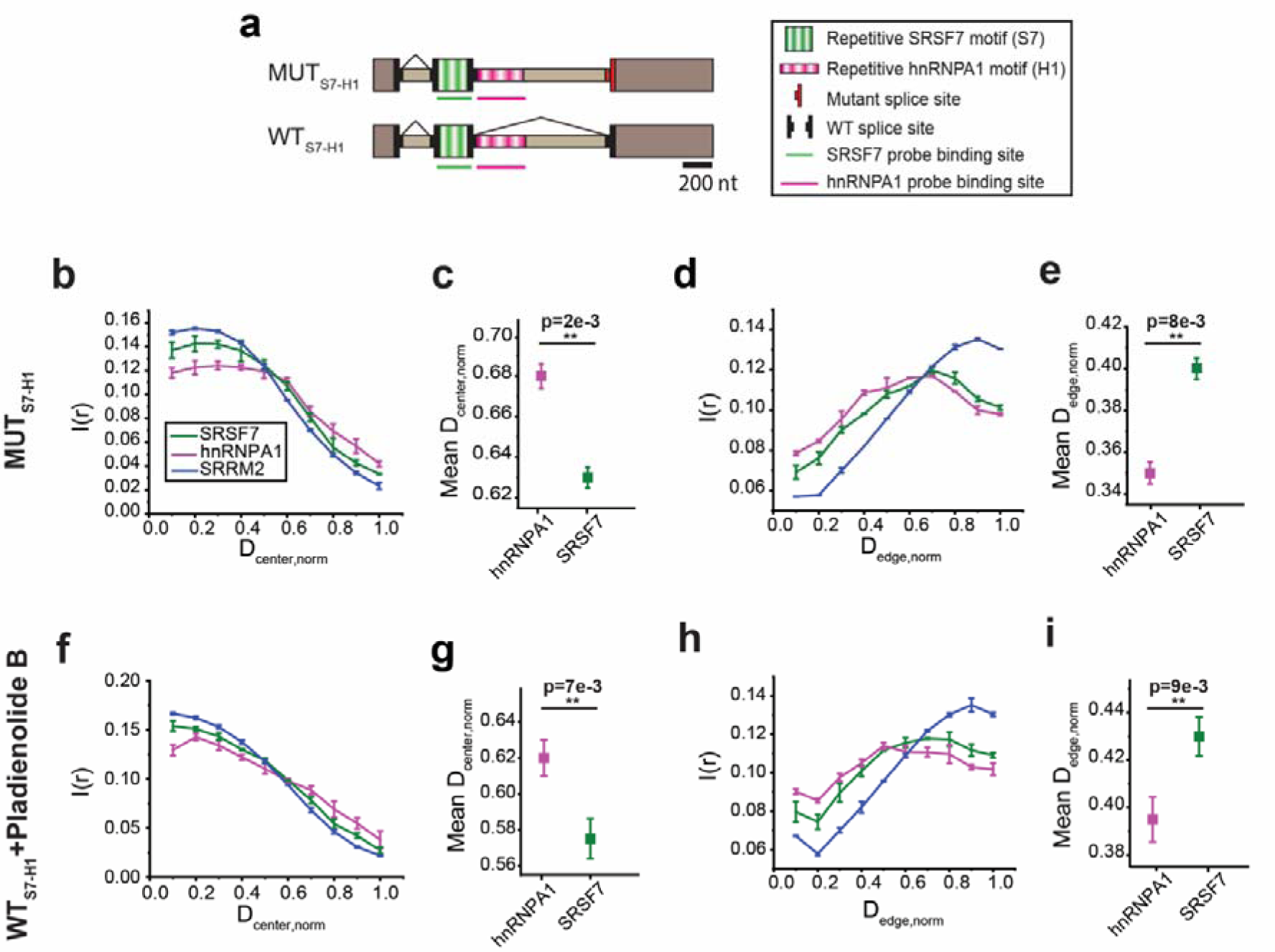
Intra-speckle organization of RNAs containing SRSF7 motifs in exon and hnRNPA1 motifs in intron. (a) Schematic illustration of MUTS7-H1 and WTS7-H1 constructs. Population distribution of SRSF7 and hnRNPA1 motif signals for MUTS7-H1 as a function of the normalized distance from the center of the speckle (b) and edge of the speckle (d). Population-weighted mean normalized distance of SRSF7 and hnRNPA1 signal from the center of speckle (c) and edge of speckle (e) for each speckle for MUTS7-H1. Population distribution of SRSF7 and hnRNPA1 motif signals for WTS7-H1 in the presence of Pladienolide B as a function of the normalized distance from the center of the speckle (f) and edge of the speckle (h). Population-weighted mean normalized distance of SRSF7 and hnRNPA1 signal from the center of speckle (g) and edge of speckle (i) for each speckle for WTS7-H1 in the presence of Pladienolide B. Error bars in the population vs. distance plots report the standard deviation from two replicates, each replicate containing at least 48-60 nuclear speckles collected from 4-5 cells. Scatter plots are generated by combining all nuclear speckles (96–120) from two replicates. Values in scatter plot represent mean ± standard error of mean (s.e.m). p-values in the scatter plots are calculated with paired sample Wilcoxon signed rank test (one-sided), with *p<5e-2, **p<1e-2, ***p<1e-3.

### RNA-RBP interaction strength determines RNA organization in nuclear speckle

We hypothesize that the mechanism underlying intra-speckle RNA organization is the multivalent interactions between RNAs with RBPs residing inside and outside nuclear speckles. We therefore expect that weakening RNA-SR protein interaction will lead to the migration of RNAs towards the speckle periphery, whereas weakening RNA-hnRNP protein interaction will lead to the migration of RNAs towards the speckle interior. In addition, weakened RNA-RBP interactions should lead to reduced constraints on RNA orientation, which would be reflected by a reduced difference in intra-speckle positioning of the two motif-rich regions. To test this hypothesis, we perturbed the RNA-RBP interaction in two ways: knocking down RBPs individually or removing RBP binding motifs from the RNA individually.

We first measured intra-speckle organization upon siRNA-mediated knockdown of SRSF1 or hnRNPA1 proteins. We achieved 83±5% and 57±8% knockdown efficiency of *SRSF1* and *hnRNPA1* mRNAs, respectively **(Figure S9).** We did not observe any significant change in nuclear speckle morphology upon knocking down of these two proteins **(Figure S8)**. We performed SMLM imaging under these knockdown conditions, using MUT_S1-H1_ as a representative. To choose cells with efficient protein knockdown, hnRNPA1 and SRSF1 proteins were stained with their respective antibodies and imaged with a 750 nm laser. Cells showing significant reduction in immunofluorescence signal compared to cells treated with scramble siRNA were selected **(Figure 5b)**. Confirming our hypothesis, SRSF1 knockdown caused significant migration of RNA transcripts towards the speckle periphery **(Figure 5c-g)**. Conversely, hnRNPA1 knockdown caused RNA migration towards the speckle interior **(Figure 5c-g)**. In addition, as predicted, the difference in intra-speckle positioning of the two motif-rich regions was reduced under each knockdown conditions. Finally, the trends were reproducible when we reversed the FISH labelling scheme **(Figure S6e-l)**.

**Figure 5.**
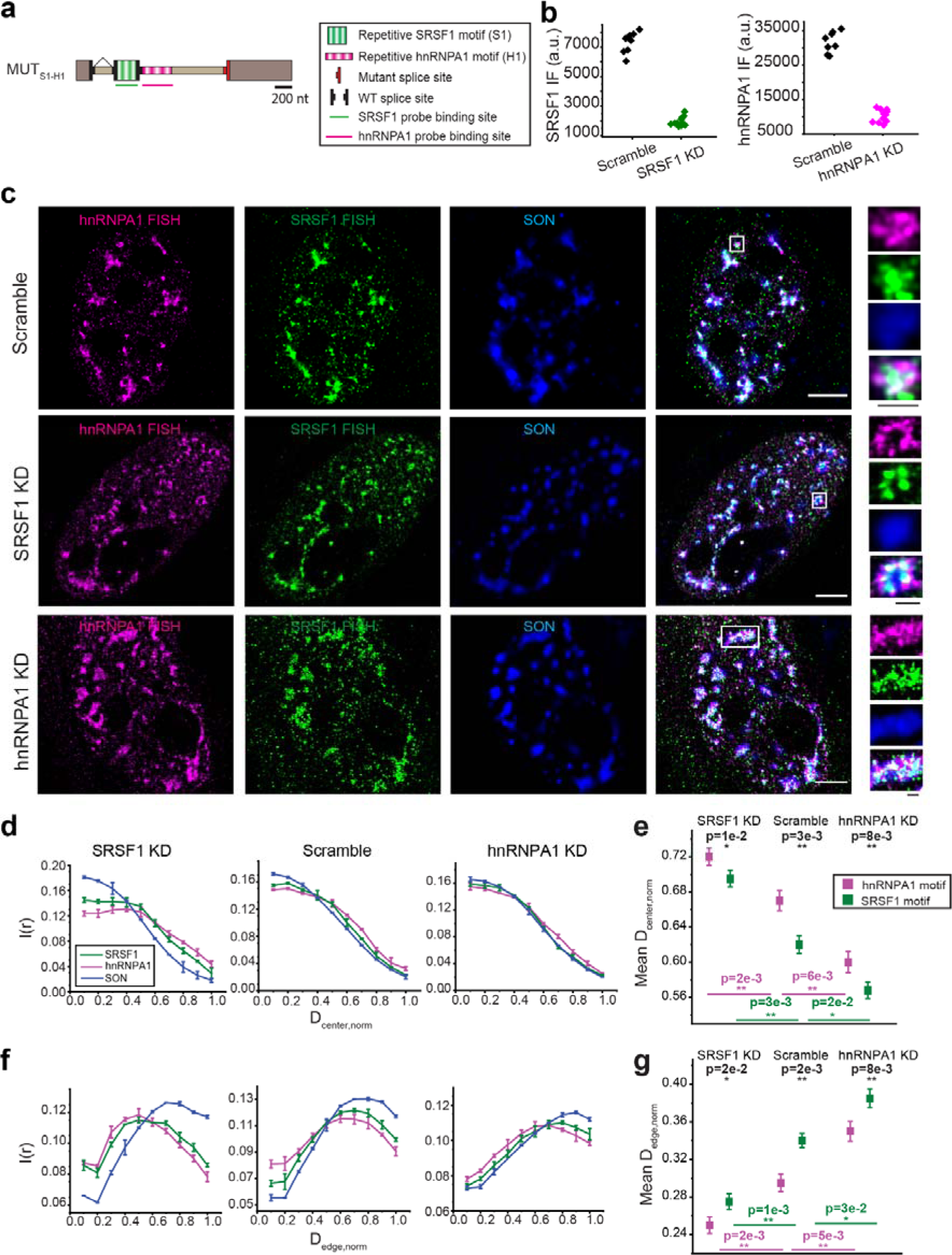
Effect of SRSF1 and hnRNPA1 knockdown on the intra-speckle organization of RNAs containing SRSF1 motifs in exon and hnRNPA1 motifs in intron. (a) Schematic illustration of MUT_S1-H1_. (b) Immunofluorescence (IF) quantification of each cell chosen for imaging treated with scramble siRNA, siRNA against SRSF1 and siRNA against hnRNPA1. Immunofluorescence is quantified by staining SRSF1 and hnRNPA1 proteins with their respective antibodies and imaging with a 750 nm laser under scramble and knockdown conditions and computing the average intensity of the whole cell. (c) Representative SMLM images of MUT_S1-H1_ treated with scramble siRNA, siRNA against SRSF1 and siRNA against hnRNPA1. FISH signals corresponding to hnRNPA1 and SRSF1 motifs in the RNAs are shown in magenta and green respectively. Immunostaining of scaffold protein SON is shown in blue. Scale bars represent 5 μm (white) and 1 μm (black). Population distribution of SRSF1 and hnRNPA1 motif signals as a function of the normalized distance from the center (d), and from the edge (f) of the speckle for MUT_S1-H1_. Population-weighted mean normalized distance of SRSF1 and hnRNPA1 signal from the center (e), and from the edge (g) for each speckle for MUT_S1-H1_. Error bars in the population vs. distance plots report the standard deviation from two replicates, each replicate containing at least 48-72 nuclear speckles collected from 4-6 cells. Scatter plots are generated by combining all nuclear speckles (96–144) from two replicates. Values in scatter plot represent mean ± standard error of mean (s.e.m). p-values in the scatter plots are calculated with paired sample Wilcoxon signed rank test (black, one-sided) and two sample t-test (magenta and green, one-sided), with *p<5e-2, **p<1e-2, ***p<1e-3.

We extended our siRNA-mediated knockdown experiments to MUT_S7-H1_. In contrast to MUT_S1-H1_, SRSF1 knockdown only caused minor outward movement of RNA transcripts from the MUT_S7-H1_ construct, and no noticeable difference in intra-speckle positioning of the two motif-rich regions **(Figure S10)**. This is consistent with the fact that MUT_S7-H1_’s middle exon does not contain any SRSF1 motifs and should thus be less sensitive to SRSF1 protein knockdown. This minor outward migration might be explained by a slight downregulation (10±9%) of SRSF7 when knocking down SRSF1 **(Figure S9).** On the other hand, as expected, hnRNPA1 knockdown still led to very similar results to those observed with MUT_S1-H1_ (migration towards speckle center and smaller difference in positioning between the two motif-rich regions) **(Figure S10)**.

We further modulated the RNA-RBP interactions by removing one of the RBP binding motifs in the RNA. We generated two size-matching single-motif variants of construct MUT_S1-H1_: MUT_N-H1_ containing only an hnRNPA1 motif-rich region and MUT_S1-N_ containing only an SRSF1 motif-rich region **(Figure 6a)**. MUT_N-H1_ RNA transcripts lacking the SRSF1 binding motifs showed a migration towards the speckle periphery **(Figure 6b-d, g-h)**, similar to the effect of SRSF1 protein knockdown. In addition, MUT_S1-N_ RNA transcripts lacking the hnRNPA1 binding motifs showed an inward migration towards the center of the speckle **(Figure 6b, e-f, g-h)**, similar to the effect of hnRNPA1 protein knockdown.

**Figure 6.**
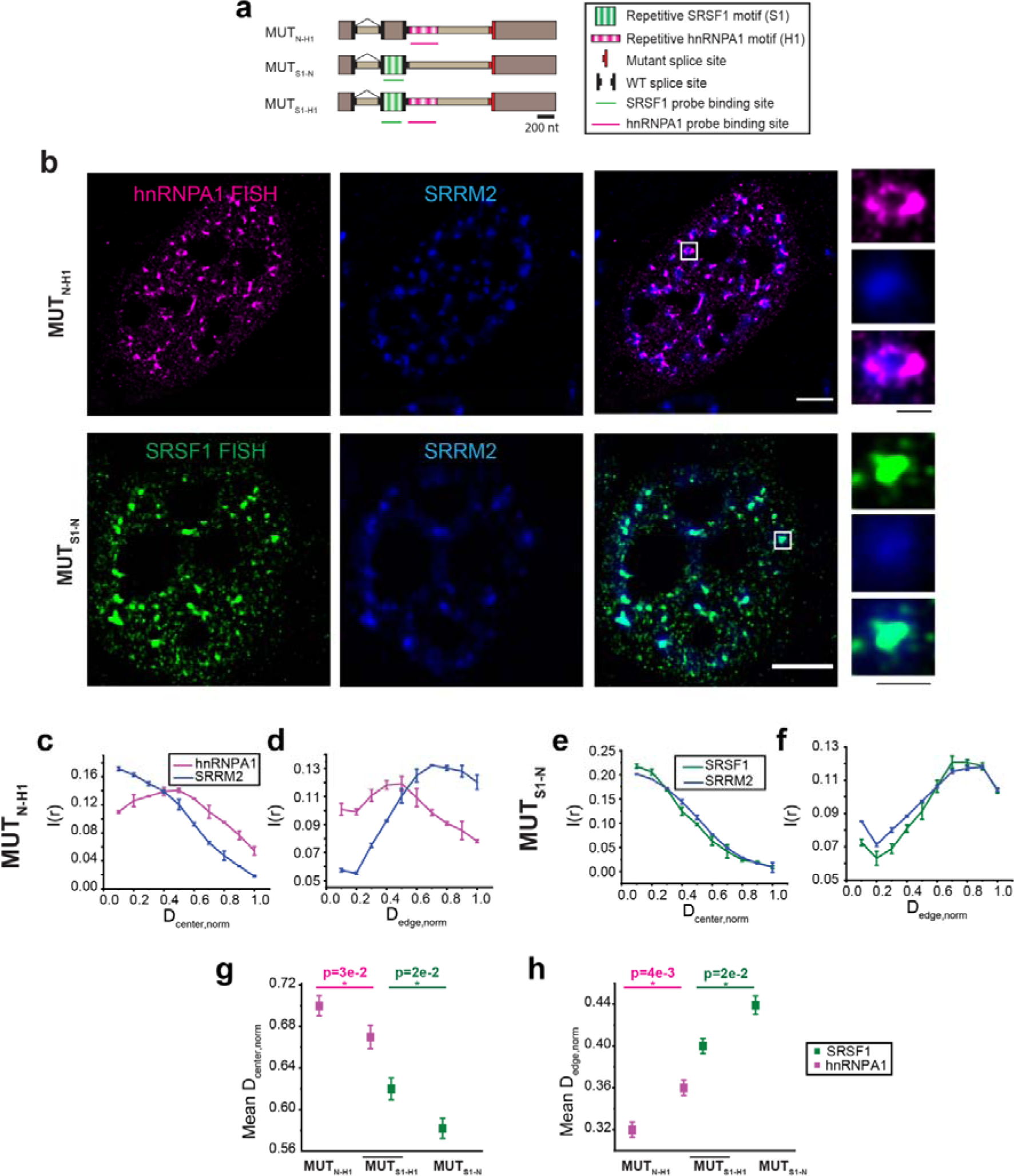
Intra-speckle organization of RNAs from single-motif constructs containing either SRSF1 motifs in exon or hnRNPA1 motifs in intron. (a) Schematic illustration of MUTN-H1, MUTS1-N and MUTS1-H1 constructs. MUTN-H1 and MUTS1-N are size-matched variants of construct MUTS1-H1. MUTN-H1 contains hnRNPA1 binding motifs in the intron and a neutral exon. MUTS1-N contains SRSF1 binding motifs in the exon and a neutral intron (b) Representative SMLM image of MUTN-H1 and MUTS1-N. Scale bars represent 5 μm (white) and 1 μm (black). Population distribution of hnRNPA1 motif signals for MUTN-H1 as a function of the normalized distance from the center of the speckle (c) and edge of the speckle (d). Population distribution of SRSF1 motif signals for MUTS1-N as a function of the normalized distance from the center of the speckle (e) and edge of the speckle (f). Comparison of the population-weighted mean normalized distance of hnRNPA1 and SRSF1 signal from MUTS1-H1 with hnRNPA1 signal from MUTN-H1 and SRSF1 signal from MUTS1-N, respectively from the center of speckle (g) and edge of speckle (h) for each speckle. Error bars in the population vs. distance plots report the standard deviation from three replicates, each replicate containing at least 75-90 nuclear speckles from 5-6 cells. Scatter plots are generated by combining all nuclear speckles (150–180) from two replicates. Values in scatter plot represent mean ± standard error of mean (s.e.m). p-values in the scatter plots are calculated with two sample t-test, with *p<5e-2, **p<1e-2, ***p<1e-3.

Collectively, these results suggest that RNA-RBP interaction strength drives intra-speckle RNA organization.

### A toy model reproduces the intra-speckle RNA organization

Finally, we computationally tested whether RNA-RBP interactions are sufficient for explaining the observed intra-speckle RNA organization. In this simple toy model, we considered four lattice sites, with two sites inside the speckle and two sites outside. We then considered the position and orientation of a 2-block RNA molecule, corresponding to an SR motif-rich region and an hnRNP motif-rich region, leading to a total of 6 configurations **(Figure 7a)**. In each configuration, each block on the RNA molecule can be either bound or unbound by the corresponding protein, leading to 4 binding states per configuration, and 24 energy states in total **(Table S1)**. The relative population of each energy state can be estimated by the dissociation constant (K_d_) of each RNA-RBP pair, and the concentration of these RBPs in each location. The intra-speckle positioning of the two regions on the RNA was then estimated using a Boltzmann distribution. The modeling details are described in Supplementary Text.

**Figure 7.**
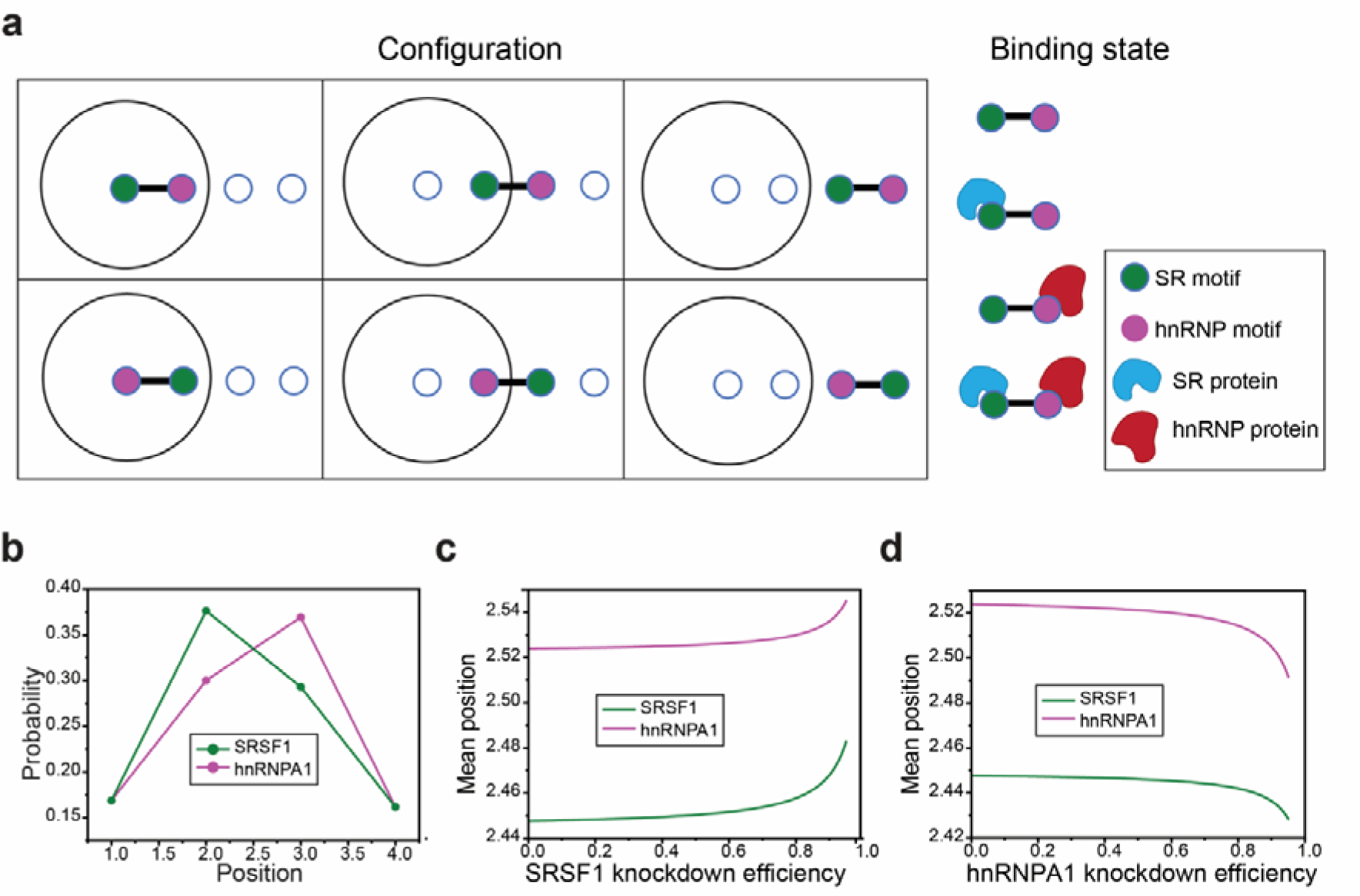
A toy model recapitulated the intra-speckle positioning of SRSF1 and hnRNPA1 motif-rich region of the RNA. (a) Graphical representation of the 6 configurations (3 positions and 2 orientations) of the RNA molecule and 4 binding states. Specifically, the positions of the SRSF1 and hnRNPA1 motif-rich regions can be both inside, both outside, or straddle the speckle interface (one inside and one outside). Each position can have two orientations (SRSF1 motif-rich region inside or hnRNPA1 motif-rich region inside). In each configuration, there are four binding states corresponding to each block on the RNA molecule being either bound or unbound by the corresponding protein. (b) Probability distribution of position of the SRSF1 and hnRNPA1 motif-rich regions, as predicted by the toy model. Mean positions of SRSF1 and hnRNPA1 motif determined by this model are plotted as a function of both SRSF1 (c) and hnRNPA1 (d) knockdown efficiencies. The case here represents Kd=1 μM.

Our simulation recapitulated the differential intra-speckle spatial distribution of the SRSF1 and hnRNPA1 motif-rich regions **(Figure 7b)**. In addition, the model recapitulated the changes of intra-speckle RNA organization upon SRSF1 and hnRNPA1 knockdowns **(Figure 7c-d)** in two aspects: (1) a change in RNA positioning as an entity, (2) a less constrained RNA orientation reflected by a smaller difference in the intra-speckle positioning of the two RBP motifs. Finally, the simulation indicated that the intra-organelle organization applies to a broad range of binding affinities (with K_d_ ranging from 100 nM to 10 µM) **(Figure 7 and S11a-f)**. In fact, increasing the K_d_ parameter in our toy model leads to results closer to the experimentally measured values, suggesting that RNA-RBP interactions in vivo may be considerably weaker than the in vitro measured values (*43*, *44*), possibly due to competition from other cellular proteins.

## Discussion

In this study, we proposed an intra-organelle RNA organization model, and demonstrated it using nuclear speckles as a model system **(Figure 1)**. Specifically, we observed that an SR motif-rich region is localized closer to the speckle center than an hnRNP motif-rich region present in the same RNA molecule. To experimentally demonstrate the spatial organization of RNA molecules, we engineered reporter constructs that are enriched in SRSF1/7 and hnRNPA1 binding motifs. Such a design guarantees that the intra-speckle organization is primarily driven by interactions with these specific proteins, and allowed us to measure the effect of knockdowns. We expect the organization of endogenous RNA transcripts to be determined by the combined interaction with the multitude of RBPs inside and outside speckles.

Intra-organelle RNA organization is likely not restricted to nuclear speckles, and can potentially apply to any membraneless organelle enriched in certain RBPs but depleted for others. Indeed, our toy model shows that RNA organization can arise from an RBP concentration difference between the inside and outside of membraneless organelles. In addition, it was reported that a long mRNA, *AHNAK* (>18 kb), that localizes to cytoplasmic stress granules, is more often observed with its 3’ end extending out of stress granules and its 5’ end residing in stress granules, compared to the other orientation (*15*). While no mechanism for this observation was provided, this observation suggests that non-random RNA organization within other membraneless organelles is possible, and remains to be further investigated.

While nuclear speckles are not known to rely on any specific RNA to assemble, their structural stability depends on the presence of nuclear RNA in general. Indeed, a recent study showed that depletion of nuclear RNA leads to loss of nuclear speckles and causes SON and SRRM2 to reorganize into a few large protein aggregates (*11*). Our observation that RNAs exhibit preferential intra-speckle organization might explain the importance of RNA molecules in maintaining the structural integrity of nuclear speckles. Specifically, through their interactions with proteins inside and outside of nuclear speckles, RNA molecules might be oriented similarly to amphiphilic polymers at the water-oil interface, and help prevent the formation of SON or SRRM2 protein aggregates. A related idea was recently proposed in the context of proteins, where the MEG-3 protein was suggested to serve as a Pickering agent to maintain an appropriate size distribution of P granules in C. elegans, by localizing to their surface and reducing surface tension (*45*).

Finally, our results suggest that nuclear speckles might play a role in facilitating splicing through organizing pre-mRNA substrates. SR and hnRNP proteins are important splicing regulators (*46*, *47*) showing antagonistic effects on splicing (*48*). While SR binding motifs are more enriched in exons, hnRNP binding motifs are more enriched in introns (*34*). We hypothesize that this specific sequence arrangement can enhance splicing by driving intra-speckle organization of pre-mRNA substrates (*49*). Specifically, a splice site found between an SR-motif rich exon and an hnRNP-motif rich intron will be positioned at the speckle outer layer, possibly providing better spatial overlap with spliceosomal components, which are known to also localize at the outer layer (*16*), thereby favoring the splicing reaction. However, future experiments are needed to demonstrate a causal relationship between intra-speckle RNA organization and splicing activity.

## Star Methods

### Plasmid design and construction

Plasmid design was based on our earlier work (*33*, *50*). Briefly, a three-exon construct was used, consisting of a Tet-responsive promoter, Chinese hamster *DHFR* exon 1 and intron 1, a synthetic variable region around exon 2, an intronic sequence derived from *DHFR* intron 3 (with a mutation in the 3’ splice site for the MUT family of constructs), and finally, the concatenation of *DHFR* exons 4 through 6 followed by the SV40 polyA sequence in the strong orientation. The SR motif-rich region in exon 2 consists of 15 repeats of an 8nt SR-binding sequence (either SRSF1 or SRSF7) separated by an 8nt “neutral” reference sequence (in total 248nt), and is flanked by 3’ and 5’ splice sites. An hnRNPA1 motif-rich region with 24 repeats was designed similarly (392nt), with the 8nt hnRNPA1 binding sequence chosen for affinity and specificity based on RNAcompete data (*51*). This region was placed downstream of exon 2 (except in WT_H1-S1_, where it was placed upstream). The neutral sequences in MUT_S1-spacer-H1,_ MUT_S1-N_, and MUT_N-H1_ were derived from repeats of the reference sequence, and were of length 720nt in MUT_S1-spacer-H1_ and size-matched in the other two cases (392nt and 248nt, respectively).

Constructs MUT_S1-H1_, MUT_S1-spacer-H1_, WT_S1-H1_, WT_H1-S1_, MUT_S1-N_, MUT_N-H1_, WT_S7-H1_, and MUT_S7-H1_ were generated in three steps. In the first step, SR motif-rich region, neutral reference sequence, and hnRNPA1 motif-rich region were generated separately producing intermediate plasmids (*33*), by following a previously published PCR-free cloning approach (*52*). Briefly, type IIS enzymes (BsaI, New England Biolabs #R3733, and BsmBI, New England Biolabs #R0739) were used to iteratively concatenate sequence modules. In the second step, the intermediate plasmids were combined with a plasmid containing the 5’ splice site or the 3’ splice site using the same stepwise approach. For example, for plasmid WT_S1-H1_, an intermediate plasmid was obtained containing the SR motif-rich region followed by a 5’ splice site and then the hnRNP motif-rich region. In the third step, the assembled sequences were transferred to the appropriate target plasmids (WT, MUT) by using a different set of type IIS enzymes (BbvI, New England Biolabs #R0173, and BfuAI, New England Biolabs #R0701). Importantly, these plasmids contain a tetracycline responsive promoter (*33*, *53*).

The non-repeat construct MUT_S1-H1,NR_ was obtained from MUT_S1-H1_ by mutating each nucleotide in the SR and hnRNP motif-rich regions with a probability of 37.5%. Doing so is likely to introduce unwanted splice site sequences, and to abolish too many of the RBP binding motifs. We therefore ranked *in silico* 10,000 candidate sequences for each region (SR rich and hnRNP rich) and picked sequences that (1) contain no predicted splice site sequence (*54*, *55*), and (2) keep a similar enrichment of SR or hnRNP motifs as scored using our previous machine learning model (*56*). The final sequences contained over 30% mutations compared to the original ones. Using BLAST of the sequence against itself, we verified the absence of any residual repeats. Gene fragments were then synthesized using gBlocks™ (IDT, USA) and cloned into the same target plasmids.

All plasmids were verified using whole-plasmid sequencing (Plasmidsaurus).

### Cell culture, transfection, and drug treatments

HeLa Tet-On cells (TaKaRa) were cultured in high glucose (4.5 g/L) containing Dulbecco’s Modified Eagle Medium (DMEM, Gibco) supplemented with 10% fetal bovine serum (FBS, Gibco), 1 mM sodium pyruvate (Gibco), 50 U/mL penicillin-streptomycin (Gibco). Cells were grown at 37 °C in a humidified environment containing 5% CO_2_. For imaging, cells were seeded in an eight-well imaging chamber (#1.5 cover glass, Cellvis) and grown overnight to 70-80% confluency before transfection.

For transfecting each well, 0.6 μL of Lipofectamine 3000 reagent (Invitrogen) was diluted in pre-warmed 15 μL reduced serum minimum essential medium (opti-MEM, Gibco) and vortexed briefly. In another tube, 200 ng of plasmid DNA and 0.4 μL P3000 reagent (Invitrogen) were diluted in 15 μL pre warmed opti-MEM and vortexed briefly. The two solutions were mixed, vortexed briefly and incubated for 15 min at room temperature. The cell culture medium was replaced with pre-warmed DMEM containing 10% Tet system approved FBS (Tet-free medium, TaKaRa) and 30 μL of DNA-lipid complex was added to each well. The medium was replaced with fresh Tet-free medium 6-8 h after transfection and incubated overnight.

Transcription induction of the transfected construct was done 24 h after transfection using 2 μg/mL doxycycline (Santa Cruz # sc-204734B) in Tet-free medium. For samples where splicing inhibition was required, cells were treated with 100 nM Pladienolide B (Plad B, Cayman) in Tet-free medium for 4 h. For transcription inhibition, induction with doxycycline was done for 2 h followed by treatment with Triptolide (40 µM, Sigma Aldrich) for 1 h.

### RT-PCR for biochemical assays

RNA was extracted 20 h after transfection using QIAgen RNeasy minikits (#74104) in a QIAcube following the manufacturer’s protocol. DNA was removed using TURBO DNase (Thermofisher #AM2238) in a 30 uL reaction. RNA was then quantified using a Nano Drop One (Thermofisher #ND-ONE-W) and the concentration adjusted to 60 ng/uL. For reverse transcription, 200 ng of RNA were used for a 10 uL reaction using SuperScript IV (Thermofisher #18090010) according to the manufacturer’s protocol. Appropriate RT primers (**Table S2**) were added at a final concentration of 100 nM. PCR reactions were carried out in a Veriti 96-well thermocycler (Applied Biosystems #4375305) using a Phusion High Fidelity kit (New England Biolabs # E0553L). The reverse transcription product was diluted 5-fold with water and 2 uL were used in a 25 uL PCR reaction according to the manufacturer’s instructions. PCR was run for 21 or 22 cycles allowing 2 min for extension. The PCR product was run in 1.5 % agarose gels and quantified in a BioRad gel documentation system after post-staining with Ethidium Bromide and destaining.

### siRNA-mediated knockdown

SRSF1 and hnRNPA1 knockdown was performed using double-stranded siRNAs against SRSF1 (hs.Ri.SRSF1.13.2, IDT, USA) and hnRNPA1 (hs.Ri.HNRNPA1.13.2, IDT, USA). A scrambled double-stranded siRNA (DsiRNA, IDT, USA) was used as negative control. Cells were seeded in an eight-well imaging chamber and grown to 60-70% confluency. For each well, 1.5 μL Lipofectamine RNAiMax reagent (Invitrogen) was diluted in 25 μL pre-warmed opti-MEM and vortexed briefly. In a separate tube, 0.5 μL siRNA (10 μM) was diluted in 25 μL pre-warmed opti-MEM and vortexed. The two solutions were then mixed, vortexed and incubated at room temperature for 5 min. 25 uL was then added to each well after replacing the cell culture medium with Tet-free medium. 6-8 h after transfection, fresh Tet-free medium was added. For the MALAT1 experiments, two rounds of siRNA mediated hnRNPA1 knockdown were done with a 24 h interval. For the experiments with constructs, single siRNA knockdown was done using Lipofectamine RNAiMax reagent followed by plasmid transfection using Lipofectamine 3000 with an interval of 24 h.

### qPCR quantification for knockdown efficiency

HeLa Tet-On cells were grown in a 12-well plate and siRNA mediated knockdown was performed. Cells were collected 48 h after knockdown first knockdown. RNA extraction was done using the Qiagen RNeasy kit (Qiagen, #75144) following the provided protocol. cDNA was synthesized using an iScript cDNA synthesis kit (BioRad). 0.5-1 μg RNA template was used and the reaction was performed in a thermal cycler (Applied Biosystems) as follows: priming for 5 min at 25 °C, reverse transcription (RT) for 20 min at 46 °C, RT inactivation for 1 min at 95 °C and then held at 4 °C. For qPCR, 2 μL of cDNA was mixed with 2 μL of forward and reverse primers (2.5 μM each, **Table S2**) and 1x SYBR Green Supermix (BioRad) for a final reaction volume of 20 μL. The qPCR reactions were performed using the CFX real-time PCR system (Bio-Rad) as follows: pre-incubation of 95 °C for 30 s, followed by 40 cycles consisting of 95 °C for 10 s and 60 °C for 30 s. The reactions were then subjected to melting curve analysis: 95 °C for 10 s, 65 °C for 5 s followed by 0.5 °C increments to 95 °C for 5 s. The data was analysed with the BioRad CFX Maestro software.

### Labelling of FISH probes and secondary antibodies

FISH probes were designed using the Stellaris Probe Designer and purchased from IDT, USA with masking level set to 5 to avoid non-specific binding. Probes were 18-20 nucleotides long with a GC content between 45%-55%. The probes targeting motifs with SRSF1, SRSF7 and hnRNPA1 in WT_S1-H1_, MUT_S1-H1_, WT_H1-S1_, WT_S7-H1_, MUT_S7-H1_, MUT_S1-N_, and MUT_N-H1_ were purchased with 3’ amine modification. For the other probes, amine modification was added using terminal transferase (TdT) enzymatic reaction (*57*). For a 60 μL reaction volume, 40 μL of pooled oligonucleotides (100 μM) were mixed with 12 μL Amino-11-ddUTP (1 mM, Lumiprobe), 2.4 μL TdT (20000 U/mL, New England Biolabs, #M0315L) in 1x TdT buffer (New England Biolabs) and incubated overnight at 37 °C in a PCR thermocycler (Applied Biosystems). The modified probes were purified using a P-6 Micro Bio-Spin Column (Bio-Rad).

For fluorophore conjugation, amine modified probes were dissolved in 0.1 M sodium bicarbonate (pH 8.5). Alexa Fluor (Invitrogen) and CF568 (Sigma Aldrich)-conjugated succinimidyl ester were dissolved in 0.5-4 μL DMSO and mixed with the probe solution. The dye: probe molar ratio was 25:1 approximately (45). The labelling reaction was incubated overnight in dark at 37 °C. To quench the reaction, 1/9^th^ reaction volume of 3M sodium acetate (pH 5) was added. Labelled probes were precipitated overnight with ethanol (∼ 2.5 times the reaction volume) and then passed through a P-6 Micro Bio-Spin column to remove unconjugated free dye. The labelling efficiency of all probes was above 75%. The exact sequences of the FISH probes are provided in Supplementary **Table S2**.

Secondary antibodies against mouse (Jackson ImmunoResearch, #715-005-150) or rabbit (Jackson ImmunoResearch, #711-005-152) were labelled with Alexa Fluor succinimidyl ester. 24 μL antibody (1 mg/mL) was mixed with 3 μL 10x PBS and 3 μL sodium bicarbonate (1 M, pH 8.5). 0.001-0.003 mg of Alexa dye was added to the above solution and the reaction was incubated for 1 h at room temperature. Labelled antibodies were purified using a P-6 Micro Bio-Spin column equilibrated with 1x PBS. 0.8 to 2.2 dye per antibody was typically achieved.

### RNA FISH and immunostaining

RNA FISH and immunostaining were performed according to a previously published protocol (*58*). Cells were fixed with 4% paraformaldehyde (PFA, Electron Microscopy Sciences) in 1x PBS for 10 min at room temperature. Permeabilization was done with a solution containing 0.5% Triton X-100 (Thermo Scientific) and 2 mM vanadyl ribonucleoside complexes (Sigma-Aldrich, #R3380) in 1x PBS for 10 min on ice. Cells were washed 3 times with 1x PBS for at least 5 min after fixation and permeabilization. Cells were stored in 70% ethanol at 4 °C until hybridization with FISH probes. Cells were washed with 2x saline-sodium citrate (SSC) two times followed by a final wash with FISH wash solution (10% formamide (Ambion, #AM9342) in 2x SSC). 125 μL of hybridization buffer (FISH wash solution and 10% dextran sulphate (Sigma-Aldrich) containing 5 nM of each labelled probe and 10 mM dithiothreitol (Sigma-Aldrich) was added to each well of the imaging chamber. The hybridization reaction was incubated overnight at 37 °C in the dark. The following day, cells were washed with FISH was solution for 30 min at 37 °C.

To prevent dissociation of probes during immunostaining, cells were again fixed with 4% PFA for 10 min at room temperature. After washing with 1x PBS, cells were treated with blocking solution (0.1% ultrapure BSA (Invitrogen, #AM2618) in 1x PBS) for 30 min at room temperature. Solutions of primary antibodies were prepared in blocking solution using the following dilutions: mouse antibody against SRRM2 (1:2000, Sigma Aldrich, #S4045), mouse antibody against SRSF1 (1:250, Invitrogen, #32-4600), rabbit antibody against hnRNPA1 (1:100, Abcam, #ab177152), rabbit antibody against SON (1:200, Invitrogen, #PA5-54814). 125 μL of the solution was added to each well and incubated at room temperature for 1 h. Cells were washed with 1x PBS three times with 5 min incubation each time. Labelled secondary antibodies were diluted 200-fold in blocking solution, 125 μL added to each well and incubated for 1 h at room temperature. Cells were washed with 1x PBS for 3 times with at least 5 min incubation time and stored in 4x SSC at 4 °C until imaging.

### Imaging and image reconstruction

Diffraction limited epi imaging was performed using a Nikon TiE microscope with a CFI HP TIRF objective (100X, NA 1.49, Nikon), and an EMCCD (Andor, iXon Ultra 888). Imaging was performed using an imaging buffer containing Tris-HCl (50 mM, pH 8), 10% glucose, 2x SSC, glucose oxidase (0.5 mg/mL, Sigma-Aldrich) and catalase (67 μg/mL, Sigma-Aldrich). FISH signals on the RNAs were imaged using the 647 nm laser (Cobolt MLD) and 561 nm laser (Coherent Obis). The immunofluorescence signal on nuclear speckle marker proteins was imaged using a 488 nm laser (Cobolt MLD). For the knockdown experiments, a 750 nm laser (Shanghai Dream Lasers Technology) was used to look at SRSF1 or hnRNPA1 protein levels stained with Alexa Fluor 750. Images were then processed in Fiji (ImageJ) (*59*) for further analysis.

2D-SMLM was performed on the same microscope, objective and EMCCD. Fluorescent TetraSpeck beads (0.1 μm, Invitrogen) were diluted 500-fold in 1x PBS, added to each well and incubated for 15 min at room temperature. After washing with 1x PBS to remove unattached beads, the same imaging buffer (as above) with additional 100 mM β-mercaptoethanol (BME, 14.3 M, Sigma-Aldrich) was added to the imaging chamber. For two color STORM, movies were collected for the Alexa Fluor 647 and CF-568 channels sequentially using JOBS module in the NIS software. Briefly, the 647 nm (∼ 40 mW) and 561 nm laser (∼ 85 mW) were used to excite Alexa Fluor 647 and CF-568 fluorophore, respectively. A 405 nm laser (CL2000, Crystal Laser) was used for activation of fluorophores from ‘off’ to ‘on’ state. The acquisition was performed with 3 frames of 647 or 561 nm laser excitation followed by 1 frame of 405 nm laser excitation, using an exposure time of 42 ms. The 405 nm laser power was adjusted during the acquisition to maintain a reasonable density of fluorophores in the ‘on-state’. The maximum 405 nm laser power used with 647 and 561 lasers was ∼2.2 mW and ∼4 mW, respectively. A total of 15000 frames were recorded for each 647 and 561 channels. Before performing SMLM imaging on a selected cell, an epi image of the same cell with at least one bead present in the region of interest was taken for channel alignment.

SMLM image reconstruction was performed using the Thunderstorm (*60*) ImageJ plugin. For approximate localization of molecules, ‘local maximum’ method was used with the peak intensity threshold 2 times the standard deviation of the residual background. To determine sub-pixel localization of molecules, the Point Spread Function (Integrated Gaussian) method was used with fitting radius of 3 pixels (pixel size = 130 nm) and initial sigma as 1.6 pixels. The ‘connectivity’ was set to ‘8-neighborhood’. The images were then corrected for translational drift using the cross-correlation method and a bin size of 20-25. Finally, spots with xy-uncertainty more than 45 nm were filtered out. Images were then rendered with 5x magnification and lateral shifts 5.

### Data analysis

A custom MATLAB code that we previously developed (*16*) was modified for radial distribution analysis on reconstructed SMLM images. Briefly, grayscale images were created from the mean intensity of all three fluorescence channels. Nuclear speckles were identified by applying an appropriate intensity threshold on the grayscale image. Inappropriately fragmented nuclear speckles were removed from the final analysis by applying a size cutoff. Further processing was done on the 2D binary images by filling and opening binary operations to remove internal voids and shot noise. Each identified nuclear speckle was indexed in region of interest (ROI), and the geometric centroid of the mask served as the center of each speckle. Additional thresholds on 2D-area and ellipticity were applied to discard abnormally large (fused) nuclear speckles and speckles that largely deviated from a spherical shape, respectively. Ellipticity was calculated as 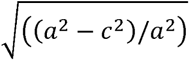, where a is the equatorial radius and c is the polar radius assuming an elliptical fit to the nuclear speckles. An area cut-off of 5000 pixels (at pixel size of 26 nm) and ellipticity cut-off of 0.8 worked best for our analysis. To estimate the ‘regularity’ of the surface of the nuclear speckles, the number of edge pixels were counted using the MATLAB built-in function *bwperim* on the generated intensity mask of the speckle and divided by the area of the speckle. For calculating the distance of the RNA motifs to the center of the nuclear speckle, the normalized radial distribution of intensity of each channel was calculated from the defined center of the speckle. The mean distance of the 647 nm and 561 nm channels (reporting the RNA signals) was calculated for each nuclear speckle, normalized by the size of the speckle (intensity weighted average radius of SRRM2/SON signal), and represented as scatter plots. For calculating the distance of the RNA motifs to the edge of the nuclear speckle, MATLAB built-in function *bwboundaries* was used to trace the exterior boundaries of nuclear speckles. For each pixel, distance to the edge is defined as the distance between that pixel and the nearest pixel on the boundary. The same procedures as described above were performed to obtain the normalized radial distribution functions and scatter plots with respect to the edge of speckles.

## Data availability

The authors declare that all data supporting the findings of the present study are available in the article and its supplementary figures and tables, or from the corresponding author upon request. Codes used for radial distribution analysis and toy model development are available on GitHub (https://github.com/JingyiFeiLab/Radial-distribution_Toy-model). All plasmids used in this study will be available through AddGene.

## Author contributions

Conceptualization: JF, OR, SP, MAA, SEL, LW Experiment and analysis: SP, MAA, LW, SEL, JZ, XW Supervision: JF, OR Writing: JF, OR, SP, LW, MAA, SEL

## References

1. M. Dundr, Nuclear bodies: multifunctional companions of the genome. Curr Opin Cell Biol. 24, 415–422 (2012).

2. L. Zhu, C. P. Brangwynne, Nuclear bodies: the emerging biophysics of nucleoplasmic phases. Curr Opin Cell Biol. 34, 23–30 (2015).

3. J. R. Buchan, R. Parker, Eukaryotic stress granules: the ins and outs of translation. Mol Cell. 36, 932–941 (2009).

4. C. J. Decker, R. Parker, P-bodies and stress granules: possible roles in the control of translation and mRNA degradation. Cold Spring Harb Perspect Biol. 4, a012286 (2012).

5. S. F. Banani, H. O. Lee, A. A. Hyman, M. K. Rosen, Biomolecular condensates: organizers of cellular biochemistry. Nat. Rev. Mol. Cell Biol. 18, 285–298 (2017).

6. S. F. Banani, A. M. Rice, W. B. Peeples, Y. Lin, S. Jain, R. Parker, M. K. Rosen, Compositional Control of Phase-Separated Cellular Bodies. Cell. 166, 651–663 (2016).

7. A. A. Hyman, C. A. Weber, F. Jülicher, Liquid-liquid phase separation in biology. Annu. Rev. Cell Dev. Biol. 30, 39–58 (2014).

8. Q. Guo, X. Shi, X. Wang, RNA and liquid-liquid phase separation. Noncoding RNA Res. 6, 92–99 (2021).

9. A. Molliex, J. Temirov, J. Lee, M. Coughlin, A. P. Kanagaraj, H. J. Kim, T. Mittag, J. P. Taylor, Phase separation by low complexity domains promotes stress granule assembly and drives pathological fibrillization. Cell. 163, 123–133 (2015).

10. Y. Lin, D. S. W. Protter, M. K. Rosen, R. Parker, Formation and Maturation of Phase-Separated Liquid Droplets by RNA-Binding Proteins. Mol. Cell. 60, 208–219 (2015).

11. C. J. Decker, J. M. Burke, P. K. Mulvaney, R. Parker, RNA is required for the integrity of multiple nuclear and cytoplasmic membrane-less RNP granules. EMBO J. 41, e110137 (2022).

12. C. M. Fare, A. Villani, L. E. Drake, J. Shorter, Higher-order organization of biomolecular condensates. Open Biol. 11, 210137 (2021).

13. J. A. West, M. Mito, S. Kurosaka, T. Takumi, C. Tanegashima, T. Chujo, K. Yanaka, R. E. Kingston, T. Hirose, C. Bond, A. Fox, S. Nakagawa, Structural, super-resolution microscopy analysis of paraspeckle nuclear body organization. J Cell Biol. 214, 817–830 (2016).

14. T. Trcek, T. E. Douglas, M. Grosch, Y. Yin, W. V. I. Eagle, E. R. Gavis, H. Shroff, E. Rothenberg, R. Lehmann, Sequence-Independent Self-Assembly of Germ Granule mRNAs into Homotypic Clusters. Mol Cell. 78, 941–950.e12 (2020).

15. S. L. Moon, T. Morisaki, A. Khong, K. Lyon, R. Parker, T. J. Stasevich, Multicolour single-molecule tracking of mRNA interactions with RNP granules. Nat Cell Biol. 21, 162–168 (2019).

16. J. Fei, M. Jadaliha, T. S. Harmon, I. T. S. Li, B. Hua, Q. Hao, A. S. Holehouse, M. Reyer, Q. Sun, S. M. Freier, R. V. Pappu, K. V. Prasanth, T. Ha, Quantitative analysis of multilayer organization of proteins and RNA in nuclear speckles at super resolution. J Cell Sci. 130, 4180–4192 (2017).

17. G. P. Faber, S. Nadav-Eliyahu, Y. Shav-Tal, Nuclear speckles - a driving force in gene expression. J Cell Sci. 135, jcs259594 (2022).

18. M. Ha, Transcription boosting by nuclear speckles. Nat Rev Mol Cell Biol. 21, 64–65 (2020).

19. J. Kim, N. C. Venkata, G. A. Hernandez Gonzalez, N. Khanna, A. S. Belmont, Gene expression amplification by nuclear speckle association. J Cell Biol. 219 (2020), doi:10.1083/jcb.201904046.

20. P. Bhat, A. Chow, B. Emert, O. Ettlin, S. A. Quinodoz, Y. Takei, W. Huang, M. R. Blanco, M. Guttman, bioRxiv, in press, doi:10.1101/2023.01.04.522632.

21. S. Hu, P. Lv, Z. Yan, B. Wen, Disruption of nuclear speckles reduces chromatin interactions in active compartments. Epigenetics Chromatin. 12, 43 (2019).

22. D. L. Spector, A. I. Lamond, Nuclear Speckles. Cold Spring Harb Perspect Biol. 3, a000646 (2011).

23. J. N. Hutchinson, A. W. Ensminger, C. M. Clemson, C. R. Lynch, J. B. Lawrence, A. Chess, A screen for nuclear transcripts identifies two linked noncoding RNAs associated with SC35 splicing domains. BMC genomics. 8, 39 (2007).

24. P. J. Shepard, K. J. Hertel, The SR protein family. Genome Biol. 10, 242 (2009).

25. A. R. Barutcu, M. Wu, U. Braunschweig, B. J. A. Dyakov, Z. Luo, K. M. Turner, T. Durbic, Z.-Y. Lin, R. J. Weatheritt, P. G. Maass, A.-C. Gingras, B. J. Blencowe, Systematic mapping of nuclear domain-associated transcripts reveals speckles and lamina as hubs of functionally distinct retained introns. Mol Cell. 82, 1035–1052.e9 (2022).

26. V. Tripathi, D. Y. Song, X. Zong, S. P. Shevtsov, S. Hearn, X.-D. Fu, M. Dundr, K. V. Prasanth, SRSF1 regulates the assembly of pre-mRNA processing factors in nuclear speckles. Mol Biol Cell. 23, 3694–3706 (2012).

27. P. J. Mintz, S. D. Patterson, A. F. Neuwald, C. S. Spahr, D. L. Spector, Purification and biochemical characterization of interchromatin granule clusters. EMBO J. 18, 4308–4320 (1999).

28. N. Saitoh, C. S. Spahr, S. D. Patterson, P. Bubulya, A. F. Neuwald, D. L. Spector, Proteomic analysis of interchromatin granule clusters. Mol. Biol. Cell. 15, 3876–3890 (2004).

29. J. Dopie, M. J. Sweredoski, A. Moradian, A. S. Belmont, Tyramide signal amplification mass spectrometry (TSA-MS) ratio identifies nuclear speckle proteins. J. Cell Biol. 219 (2020), doi:10.1083/jcb.201910207.

30. A. P. Dias, K. Dufu, H. Lei, R. Reed, A role for TREX components in the release of spliced mRNA from nuclear speckle domains. Nature Communications. 1, 97 (2010).

31. K. M. Neugebauer, M. B. Roth, Distribution of pre-mRNA splicing factors at sites of RNA polymerase II transcription. Genes Dev. 11, 1148–1159 (1997).

32. A. Abdrabou, Z. Wang, Regulation of the nuclear speckle localization and function of Rac1. FASEB J. 35, e21235 (2021).

33. M. A. Arias, A. Lubkin, L. A. Chasin, Splicing of designer exons informs a biophysical model for exon definition. RNA. 21, 213–229 (2015).

34. W. G. Fairbrother, R.-F. Yeh, P. A. Sharp, C. B. Burge, Predictive identification of exonic splicing enhancers in human genes. Science. 297, 1007–1013 (2002).

35. X.-D. Fu, M. Ares, Context-dependent control of alternative splicing by RNA-binding proteins. Nat. Rev. Genet. 15, 689–701 (2014).

36. R. Das, K. Dufu, B. Romney, M. Feldt, M. Elenko, R. Reed, Functional coupling of RNAP II transcription to spliceosome assembly. Genes Dev. 20, 1100–1109 (2006).

37. O. Gozani, J. G. Patton, R. Reed, A novel set of spliceosome-associated proteins and the essential splicing factor PSF bind stably to pre-mRNA prior to catalytic step II of the splicing reaction. EMBO J. 13, 3356–3367 (1994).

38. İ. A. Ilik, M. Malszycki, A. K. Lübke, C. Schade, D. Meierhofer, T. Aktaş, SON and SRRM2 are essential for nuclear speckle formation. Elife. 9, e60579 (2020).

39. A. Gopal, Z. H. Zhou, C. M. Knobler, W. M. Gelbart, Visualizing large RNA molecules in solution. RNA. 18, 284–299 (2012).

40. E. Lubeck, L. Cai, Single-cell systems biology by super-resolution imaging and combinatorial labeling. Nat Methods. 9, 743–748 (2012).

41. M. J. Rust, M. Bates, X. Zhuang, Sub-diffraction-limit imaging by stochastic optical reconstruction microscopy (STORM). Nat. Methods. 3, 793–795 (2006).

42. T. Carvalho, S. Martins, J. Rino, S. Marinho, M. Carmo-Fonseca, Pharmacological inhibition of the spliceosome subunit SF3b triggers exon junction complex-independent nonsense-mediated decay. J Cell Sci. 130, 1519–1531 (2017).

43. S. Cho, A. Hoang, R. Sinha, X.-Y. Zhong, X.-D. Fu, A. R. Krainer, G. Ghosh, Interaction between the RNA binding domains of Ser-Arg splicing factor 1 and U1-70K snRNP protein determines early spliceosome assembly. Proc Natl Acad Sci U S A. 108, 8233–8238 (2011).

44. C. G. Burd, G. Dreyfuss, RNA binding specificity of hnRNP A1: significance of hnRNP A1 high-affinity binding sites in pre-mRNA splicing. EMBO J. 13, 1197–1204 (1994).

45. A. W. Folkmann, A. Putnam, C. F. Lee, G. Seydoux, Regulation of biomolecular condensates by interfacial protein clusters. Science. 373, 1218–1224 (2021).

46. J. Zhu, A. Mayeda, A. R. Krainer, Exon identity established through differential antagonism between exonic splicing silencer-bound hnRNP A1 and enhancer-bound SR proteins. Mol. Cell. 8, 1351–1361 (2001).

47. K. J. Hertel, Combinatorial control of exon recognition. J. Biol. Chem. 283, 1211–1215 (2008).

48. S. Erkelenz, W. F. Mueller, M. S. Evans, A. Busch, K. Schöneweis, K. J. Hertel, H. Schaal, Position-dependent splicing activation and repression by SR and hnRNP proteins rely on common mechanisms. RNA. 19, 96–102 (2013).

49. S. E. Liao, O. Regev, Splicing at the phase-separated nuclear speckle interface: a model. Nucleic Acids Res. 49, 636–645 (2021).

50. X. H.-F. Zhang, M. A. Arias, S. Ke, L. A. Chasin, Splicing of designer exons reveals unexpected complexity in pre-mRNA splicing. RNA. 15, 367–376 (2009).

51. D. Ray, H. Kazan, E. T. Chan, L. Peña Castillo, S. Chaudhry, S. Talukder, B. J. Blencowe, Q. Morris, T. R. Hughes, Rapid and systematic analysis of the RNA recognition specificities of RNA-binding proteins. Nat Biotechnol. 27, 667–670 (2009).

52. A. Scior, S. Preissler, M. Koch, E. Deuerling, Directed PCR-free engineering of highly repetitive DNA sequences. BMC Biotechnol. 11, 87 (2011).

53. M. Gossen, H. Bujard, Tight control of gene expression in mammalian cells by tetracycline-responsive promoters. Proc Natl Acad Sci U S A. 89, 5547–5551 (1992).

54. G. Yeo, C. B. Burge, Maximum entropy modeling of short sequence motifs with applications to RNA splicing signals. J Comput Biol. 11, 377–394 (2004).

55. M. S. Wong, J. B. Kinney, A. R. Krainer, Quantitative Activity Profile and Context Dependence of All Human 5’ Splice Sites. Mol Cell. 71, 1012–1026.e3 (2018).

56. S. E. Liao, M. Sudarshan, O. Regev, bioRxiv, in press, doi:10.1101/2022.10.01.510472.

57. I. Gaspar, F. Wippich, A. Ephrussi, Enzymatic production of single-molecule FISH and RNA capture probes. RNA. 23, 1582–1591 (2017).

58. A. Raj, P. van den Bogaard, S. A. Rifkin, A. van Oudenaarden, S. Tyagi, Imaging individual mRNA molecules using multiple singly labeled probes. Nat. Methods. 5, 877–879 (2008).

59. J. Schindelin, I. Arganda-Carreras, E. Frise, V. Kaynig, M. Longair, T. Pietzsch, S. Preibisch, C. Rueden, S. Saalfeld, B. Schmid, J.-Y. Tinevez, D. J. White, V. Hartenstein, K. Eliceiri, P. Tomancak, A. Cardona, Fiji: an open-source platform for biological-image analysis. Nat Methods. 9, 676–682 (2012).

60. M. Ovesný, P. Křížek, J. Borkovec, Z. Svindrych, G. M. Hagen, ThunderSTORM: a comprehensive ImageJ plug-in for PALM and STORM data analysis and super-resolution imaging. Bioinformatics. 30, 2389–2390 (2014).

