## Supplementary text and figures for "RNA molecules display distinctive organization at nuclear speckles"

We employed a toy model, in which we considered the space as four lattice sites with two in the nuclear speckle and two in the nucleoplasm. The RNA molecule was modeled as adjacent SRSF1 and hnRNPA1 motifs, which were allowed to occupy two adjacent lattice sites. As a result, there were 6 possible configurations of RNA positions and orientations. Furthermore, each configuration can be found in one of four binding states corresponding to SRSF1 and hnRNPA1 protein being either bound or unbound (**Figure 7a**). We denote the occupancy of SRSF1 and hnRNPA1 as $\sigma_{s}$ and $\sigma_{h}$ respectively, taking value 0 (unbound) or 1 (bound). A detailed summary of the 24 states, considering all the possible RNA spatial configurations as well as protein binding, can be found in Supplementary **Table S1**.

The energy for the $i$th state can be written as

$$E^{i}=\sigma_{s}^{i}\varepsilon_{s}^{i}+\sigma_{h}^{i}\varepsilon_{h}^{i}.$$

$\varepsilon$ is a measure of RNA-RBP interaction and is given by

$$\varepsilon=-\frac{1}{\beta}ln(\frac{c}{K_{d}})$$

where $c$ is the RBP concentration in each compartment and $K_{d}$ is the dissociation constant of the RBP to its RNA motif. Here $\varepsilon$ is not the usual RNA-RBP binding affinity defined by $\frac{1}{\beta}ln(\frac{K_{d}}{1 M})$, but an effective energy to account for the concentration dependence of binding probability. With this definition of $\varepsilon$, the probability of the protein bound state follows a Boltzmann distribution $p_{b}=\frac{c/K_{d}}{1+c/K_{d}}=\frac{e^{-\beta\varepsilon}}{1+e^{-\beta\varepsilon}}$.

The effective energy between SRSF1 and its binding motif ($\varepsilon_{s}^{i}$), and the effective energy between hnRNPA1 and its binding motif ($\varepsilon_{h}^{i}$) depend on SRSF1 and hnRNPA1 motif position, and are given by

$$\varepsilon_{s}^{i}=\left\{ \begin{aligned} \varepsilon_{s1}=-\frac{1}{\beta}ln\left( \frac{c_{s1}}{K_{d}} \right), x_{s}^{i}\leq2 \\ \varepsilon_{s2}=-\frac{1}{\beta}ln\left( \frac{c_{s2}}{K_{d}} \right), x_{s}^{i}>2 \end{aligned} \right.$$

$$\varepsilon_{h}^{i}=\left\{ \begin{aligned} \varepsilon_{h1}=-\frac{1}{\beta}ln\left( \frac{c_{h1}}{K_{d}} \right), x_{h}^{i}\leq2 \\ \varepsilon_{h2}=-\frac{1}{\beta}ln\left( \frac{c_{h2}}{K_{d}} \right), x_{h}^{i}>2 \end{aligned} \right.$$

where $\varepsilon_{s1}$ and $\varepsilon_{s2}$ denote the effective energy between SRSF1 and its binding motif in nuclear speckles and nucleoplasm, $c_{s1}$ and $c_{s2}$ denote nuclear speckle and nucleoplasmic concentration of SRSF1, $\varepsilon_{h1}$ and $\varepsilon_{h2}$ denote the effective energy between hnRNPA1 and its binding motif in nuclear speckles and nucleoplasm, $c_{h1}$ and $c_{h2}$ denote nuclear speckle and nucleoplasmic concentration of hnRNPA1, $x_{s}^{i}$ and $x_{h}^{i}$ denote SRSF1 and hnRNPA1 motif position for the $i$th state taking values in $\{1,2,3,4\}$.

To determine values for $\varepsilon_{s1}$ and $\varepsilon_{s2}$, we first estimated SRSF1 concentration in nuclear speckles ($c_{s1}$) and nucleoplasm ($c_{s2}$). They are given by $c_{s1}=N_{1}/V_{1}$ and $c_{s2}=N_{2}/V_{2}$, where $N_{1}$ and $N_{2}$ denote the number of molecules in nuclear speckles and nucleoplasm, $V_{1}$ and $V_{2}$ denote the volume of nuclear speckles and nucleoplasm. The copy number of SRSF1 proteins in HeLa cells was measured earlier as $4.4\times{10}^{6}$ molecules per cells by mass spectrometry (*1*). Because SRSF1 proteins predominantly localize to nucleus, we used this number as the number of molecules in nucleus. By immunofluorescence images, we found 17.4% of SRSF1 signals present in nuclear speckles and 82.6% present in nucleoplasm. $N_{1}$ and $N_{2}$ were therefore estimated to be $7.7\times{10}^{5}$ and $3.6\times{10}^{6}$. The average volume of HeLa cell nucleus was reported to be 220 fL (*2*). Using our imaging data, we estimated that 14.0% of nuclear space was occupied by nuclear speckles$.$ We therefore calculated $V_{1}$ and $V_{2}$ to be 30.8 fL and 189.2 fL. $c_{s1}$ and $c_{s2}$ were then estimated to be 41.0 µM and 31.7 µM, respectively. $\varepsilon_{s1}$ and $\varepsilon_{s2}$ were determined by assuming $K_{d}$ to be 1 µM.

The nucleoplasmic concentration of hnRNPA1 protein $c_{h2}$ was estimated as 36.4 µM based on reported values (*3*). To estimate the nuclear speckle concentration of hnRNPA1 $c_{h1}$, we analyzed the depletion of hnRNPA1 intensity in immunofluorescence images. We found 55% of nuclear speckles exhibited 30% depletion in hnRNPA1, 30% exhibited 6% depletion, and 15% exhibited no depletion. Based on this, we generated three separate simulations recapitulating various degrees of depletion in hnRNPA1 and computed the population average. Again, we used $K_{d}$ as 1 µM to determine $\varepsilon_{h1}$ and $\varepsilon_{h2}$.

When simulating knockdown experiments, the protein concentrations were multiplied by $1-x$, where $x$ represents the knockdown efficiency, $\varepsilon_{s1}$, $\varepsilon_{s2}$, $\varepsilon_{h1}$, and $\varepsilon_{h2}$ vary accordingly as concentrations decrease.

Given the energy $E^{i}$, we can find the partition function and the probability for the $i$th state by the Boltzmann distribution

$$Z=\sum_{i=1}^{24} e^{-\beta E_{i}} , P^{i}=\frac{e^{-\beta E_{i}}}{Z}.$$

The probability distribution of SRSF1 motif position$P\left( x_{s}=j \right)$ and hnRNPA1 motif position $P\left( x_{h}=j \right)$ are given by the sum of the probabilities for states with SRSF1 motif position equal to $j$ and hnRNPA1 motif position equal to $j$ respectively. Then the mean position of SRSF1 and hnRNPA1 are determined by

$$\bar{x_{s}}=\sum_{j=1}^{4} jP(x_{s}=j) , \bar{x_{h}}=\sum_{j=1}^{4} jP(x_{h}=j)$$

All analytical expressions for the partition function and the mean position of SRSF1 and hnRNPA1 motif were obtained by Wolfram Mathematica as

$$Z=e^{-\beta\varepsilon_{h1}}\left( 3+2e^{-\beta\varepsilon_{s1}}+e^{-\beta\varepsilon_{s2}} \right)+e^{-\beta\varepsilon_{h2}}\left( 3+e^{-\beta\varepsilon_{s1}}+2e^{-\beta\varepsilon_{s2}} \right)+6+3e^{-\beta\varepsilon_{s1}}+3e^{-\beta\varepsilon_{s2}}$$

$$\bar{x_{s}}=\frac{e^{-\beta\varepsilon_{h1}}\left( 6+3e^{-\beta\varepsilon_{s1}}+{3e}^{-\beta\varepsilon_{s2}} \right)+e^{-\beta\varepsilon_{h2}}\left( 9+{2e}^{-\beta\varepsilon_{s1}}+7e^{-\beta\varepsilon_{s2}} \right)+15+5e^{-\beta\varepsilon_{s1}}+10e^{-\beta\varepsilon_{s2}}}{Z}$$

$$\bar{x_{h}}=\frac{e^{-\beta\varepsilon_{h1}}\left( 5+3e^{-\beta\varepsilon_{s1}}+{2e}^{-\beta\varepsilon_{s2}} \right)+e^{-\beta\varepsilon_{h2}}\left( 10+{3e}^{-\beta\varepsilon_{s1}}+7e^{-\beta\varepsilon_{s2}} \right)+15+6e^{-\beta\varepsilon_{s1}}+9e^{-\beta\varepsilon_{s2}}}{Z}$$

$$\bar{x_{h}}-\bar{x_{s}}=\frac{e^{-\beta\varepsilon_{h2}}\left( 1+e^{-\beta\varepsilon_{s1}} \right)-e^{-\beta\varepsilon_{h1}}\left( 1+e^{-\beta\varepsilon_{s2}} \right)+e^{-\beta\varepsilon_{s1}}-e^{-\beta\varepsilon_{s2}}}{Z}$$

These expressions indicate that this model can account for the differential intra-speckle spatial distribution of SRSF1 and hnRNPA1 motif, which is driven by the enrichment of SRSF1 and depletion of hnRNPA1 in nuclear speckles (**Figure 7b**). In agreement with our observations, this model predicts that both SRSF1 and hnRNPA1 motifs migrate towards the speckle periphery upon SRSF1 knockdown and towards the speckle interior upon hnRNPA1 knockdown (**Figure 7c-d**). Moreover, the difference in spatial distribution of SRSF1 and hnRNPA1 motif is maintained under knockdown conditions, consistent with our experimental observations (**Figure 7c-d**).

We finally tested this model applies to a broad range of binding affinities by increasing $K_{d}$ to 10 µM and decreasing $K_{d}$ to 100 nM. Similar trends were found in **Figure S10**.

**Supplementary figures**

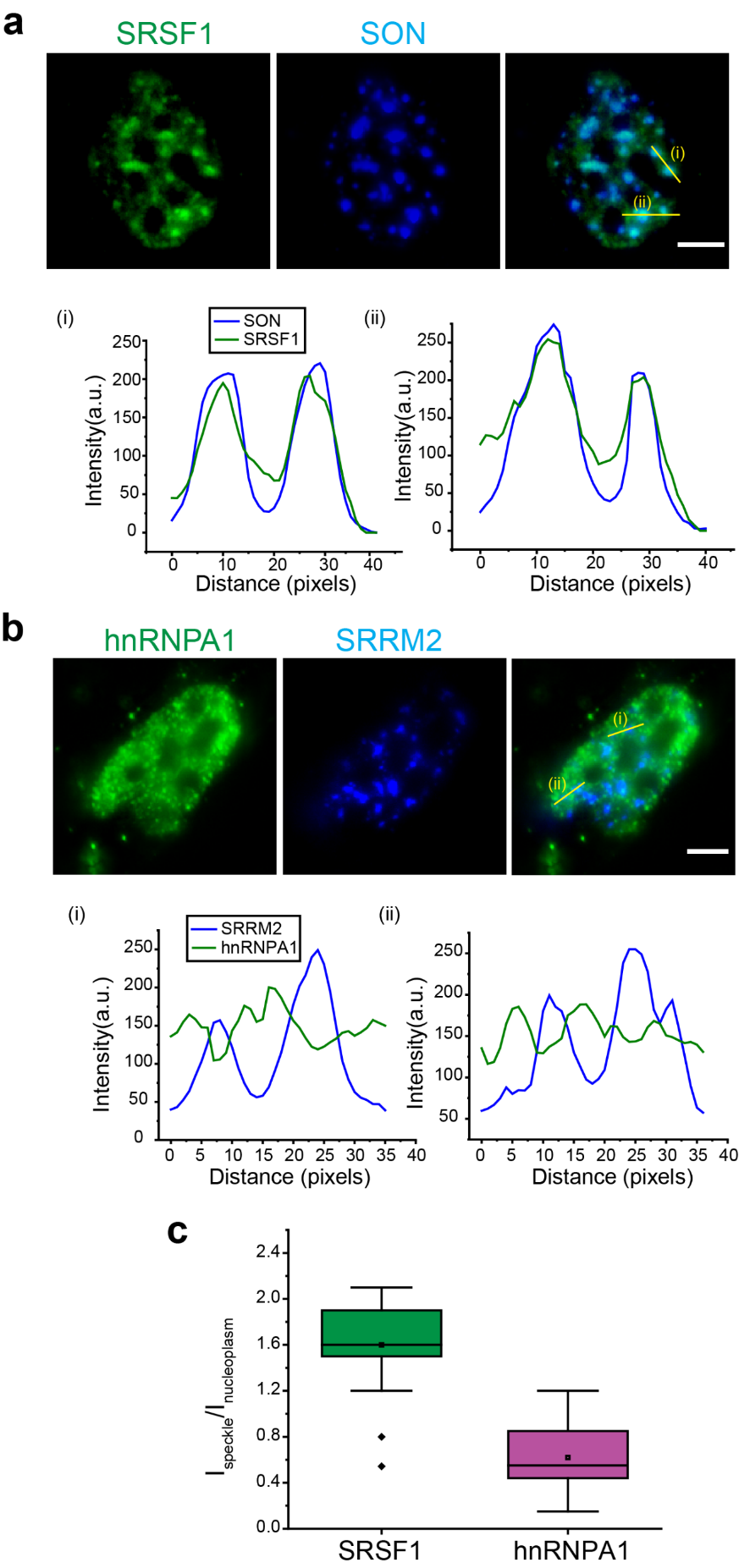

**Figure S1.** **Diffraction-limited epifluorescence images of SRSF1 and hnRNPA1 proteins.** SRSF1 (a) and hnRNPA1 (b) proteins are labeled by immunofluorescence staining using CF568, together with SON or SRRM2 protein stained with AF488. Representative speckles are highlighted to demonstrate distributions of SRSF1 and hnRNPA1 relative to nuclear speckles. (c) Comparison of SRSF1 and hnRNPA1 enrichment in nuclear speckles from ~100 speckles from 10 cells. I_speckle_ and I_nucleoplasm_ represent the mean intensities of proteins in the nuclear speckles and nucleoplasm, respectively. SRSF1 proteins are enriched in the speckles. hnRNPA1 proteins are distributed in the nucleoplasm with depletion in ~85% speckles. Scale bars represent 5 μm. Description of box plot: center line reports the median; dot inside box reports the mean; box limits are upper and lower quartiles; whiskers are 1.5x interquartile range; and points outside box are outliers.

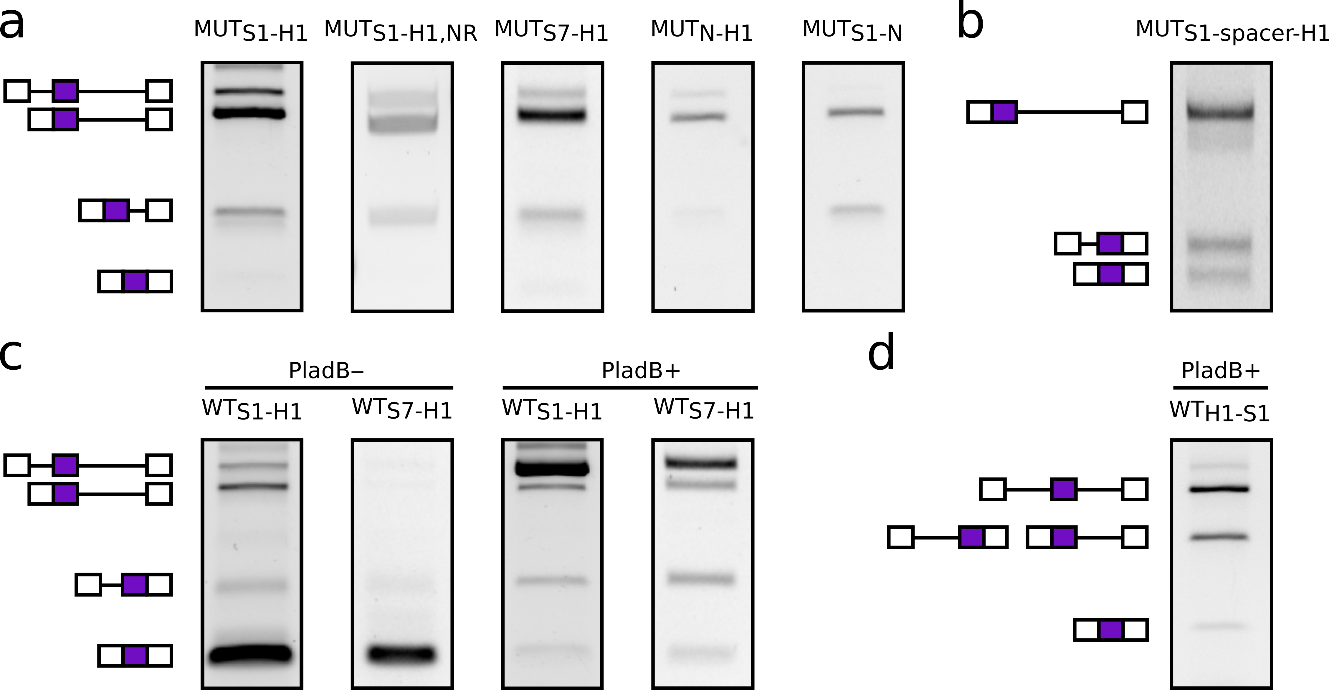

**Figure S2**. **Reverse-transcription PCR (RT-PCR) assay to test splicing outcomes.** (a) MUT_S1-H1,_ MUT_S1-H1,NR_, MUT_S7-H1_, MUT_N-H1_ and MUT_S1-N_ constructs with a 3’ splice site mutation. (b) MUT_S1-spacer-H1_ construct with a 3’ splice site mutation and a 720-nt long spacer between the SRSF1 and hnRNPA1 motif-rich regions. In all these constructs, the middle exon and its downstream intron are mostly not spliced. (c) WT_S1-H1_ and WT_S7-H1_ in the absence and presence of Pladienolide B. These constructs are spliced with the middle exon included in the absence of Pladienolide B, while in the presence of Pladienolide B, splicing is inhibited. (d) WT_H1-S1_ in the presence of Pladienolide B. Splicing is mostly inhibited in the presence of Pladienolide B. The middle band is a mixture of two products of nearly identical sizes (intron 1 retained and intron 2 retained). Induction time for all constructs is 2h.

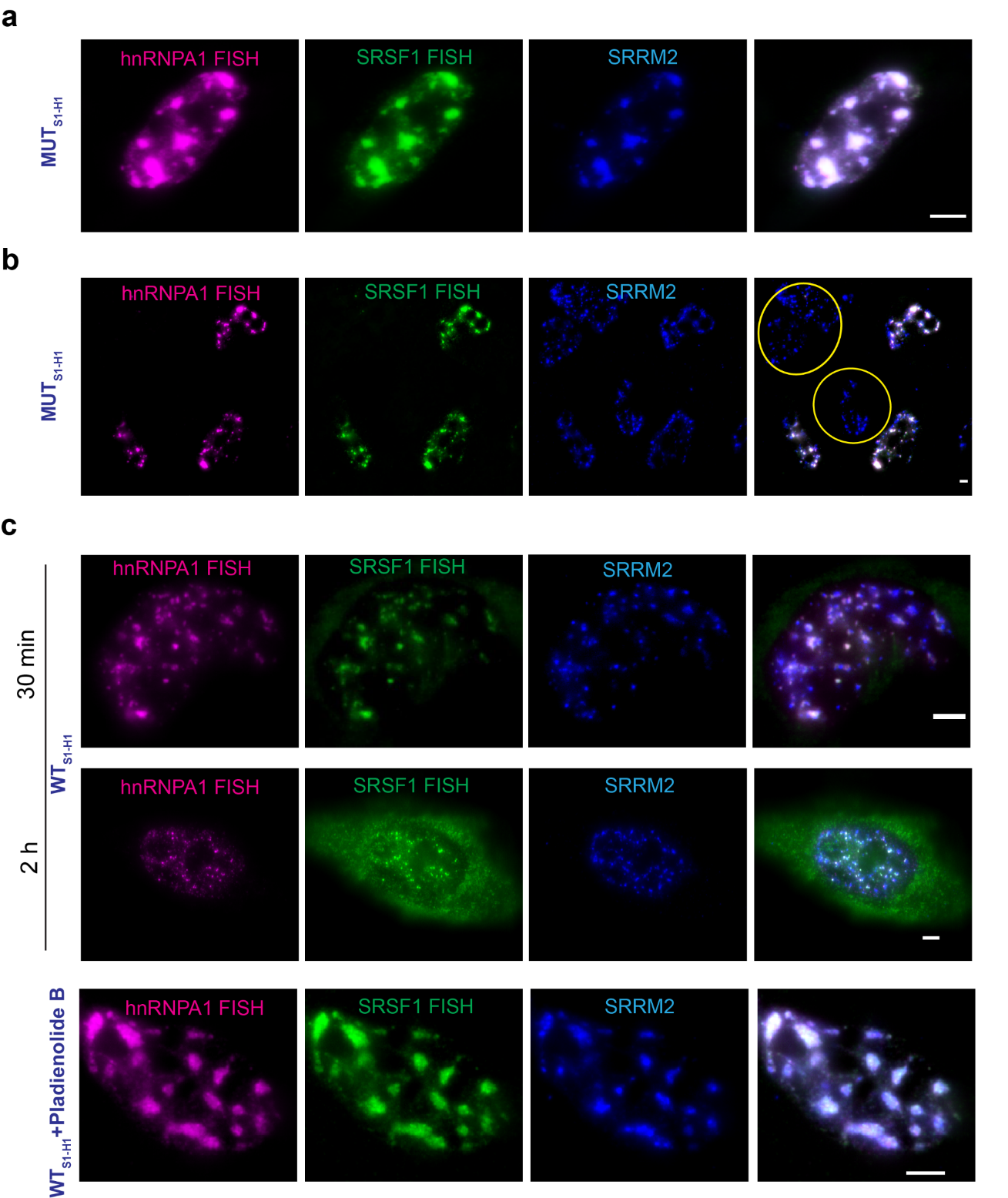

**Figure S3.** **Representative epifluorescence images of RNAs expressed from the reporter constructs.** FISH signals corresponding to hnRNPA1 and SRSF1 protein binding motifs in the RNAs are shown in magenta and green respectively. Immunostaining of SRRM2 is shown in blue. (a) 2h Induction of MUT_S1-H1_ in the absence of Pladienolide B. (b) Specificity of RNA FISH probes tested with transfected cells and cells with unsuccessful transfection as control (highlighted in yellow). The cells with unsuccessful transfection have negligible FISH signal as compared to the transfected cells. (c) 2h Induction of WT_S1-H1_ in the absence and presence of Pladienolide B. Scale bars represent 5 μm.

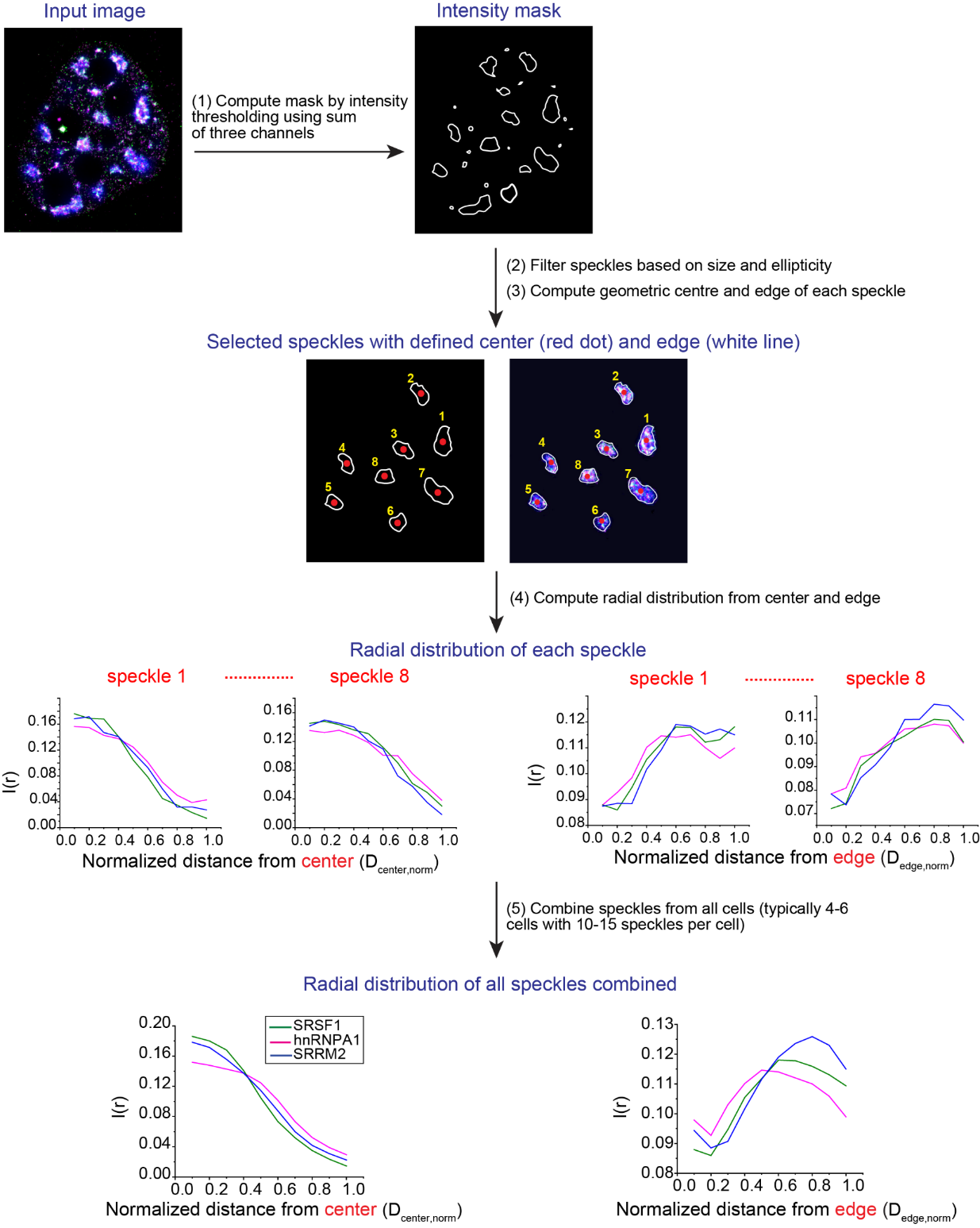

**Figure S4.** **Data analysis pipeline**. Input images are in the form of red (R), green (G) and blue (B) where R and G channels represent RNA FISH signals and B corresponds to SRRM2 immunostaining signal. A composite RGB image is generated which is then converted to a grayscale image. Using the sum of RGB channels, a mask is generated through intensity thresholding. Following this a size filter (lower limit 300 pixels, upper limit 5000 pixels) and ellipticity filter (cutoff 0.8) is applied. After defining the geometric center and edge of each selected speckle, the radial distribution is calculated for each speckle. Finally, the radial distribution of each selected speckle was averaged to generate the final plots.

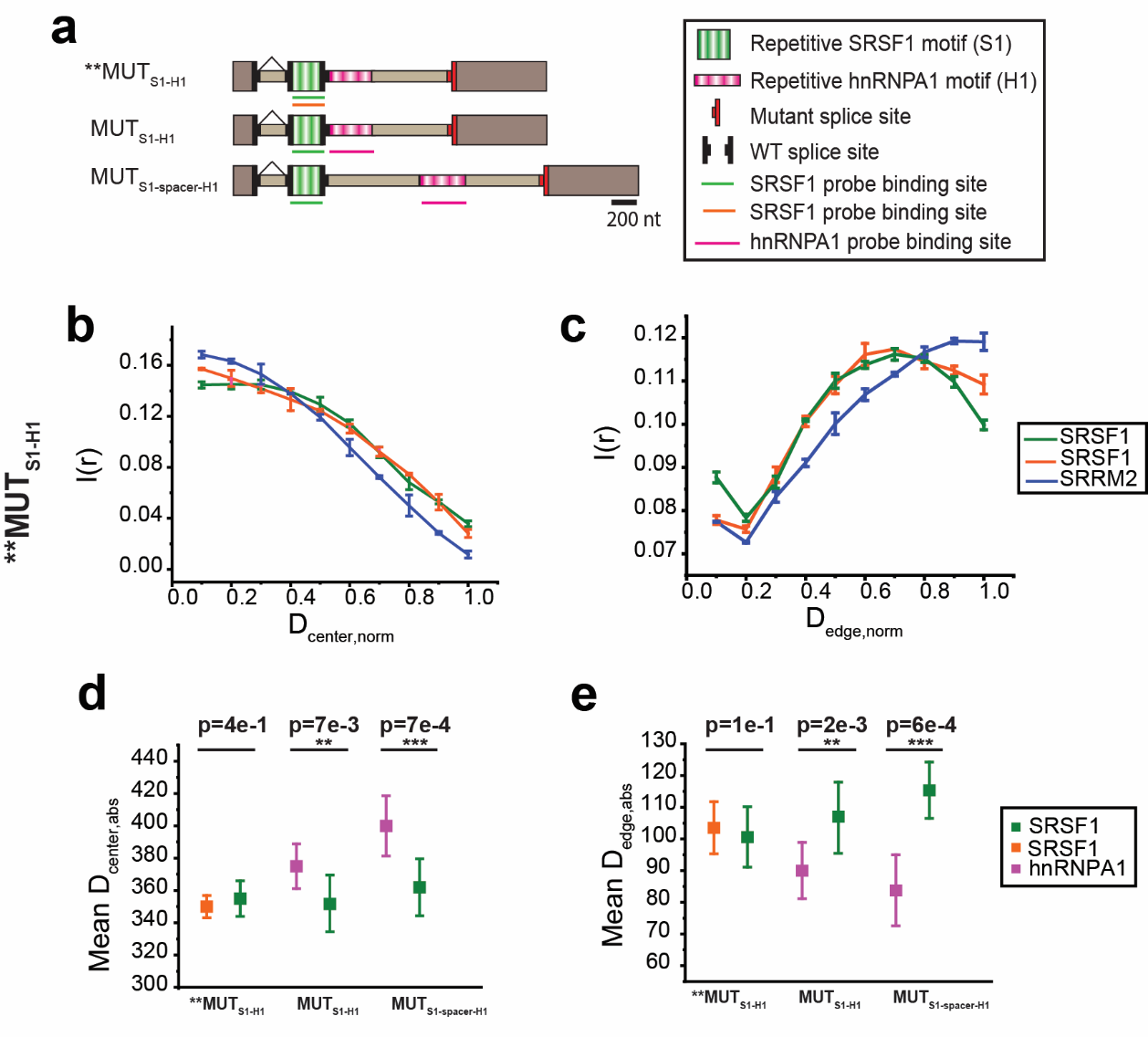

**Figure S5. Validation of mean radial distance difference.** Population distribution of SRSF1 motif signals labeled with both A647 and CF568 for MUT_S1-H1_ as a function of the normalized distance from the center of the speckle (a) and edge of the speckle (b). Population-weighted mean absolute distance of SRSF1 and hnRNPA1 signal from the center of speckle (c) and edge of the speckle (d) for each speckle for MUT_S1-H1_ and MUT_S1-spacer-H1_. Each data set contains at least 80-120 nuclear speckles collected from 8-12 cells. Values in scatter plot represent mean ± standard error of mean (s.e.m). p-values in the scatter plots are calculated with paired sample Wilcoxon signed rank test, with *p<5e-2, **p<1e-2, ***p<1e-3. ** MUT_S1-H1_ represents the case where both the A647 and CF568 labeled RNA FISH probes are on the SRSF1 motifs.

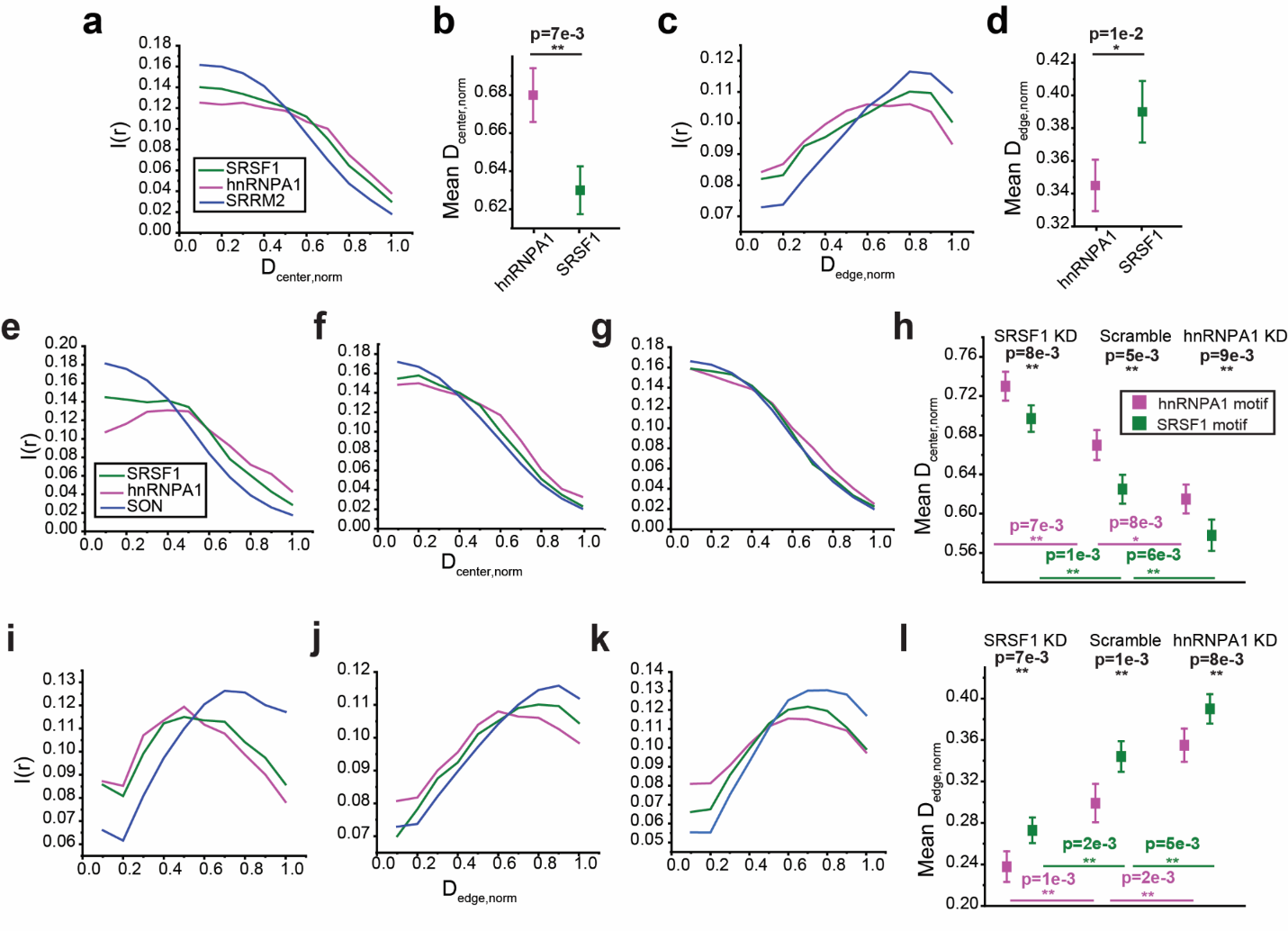

**Figure S6.** **SMLM imaging and analysis with reversed labeling of probes.** The reversed labeling scheme: hnRNPA1 motifs labeled with CF568 and SRSF1 motifs labeled with AF647. (a-d) Analysis of MUT_S1-H1_ RNA. Population distribution of SRSF1 and hnRNPA1 motif signals as a function of the normalized distance from the center (a) and from the edge (c) of the speckle. Scatter plot of the population-weighted mean normalized distance of SRSF1 and hnRNPA1 signal from the center (b) and from the edge (d) of speckle for each speckle. (e-l) Analysis of RNA positioning from MUT_S1-H1_ under knockdown conditions Population distribution of SRSF1 and hnRNPA1 motif signals as a function of the normalized distance from the center (e-g) and from the edge (i-k) of the speckle. Population-weighted mean normalized distance of SRSF1 and hnRNPA1 signal from the center (h) and from the edge (l) of speckle for each speckle. Since the reversed labeling scheme is only used for validation, data was only collected once for each case. Each data set contains 50-60 nuclear speckles collected from 5-6 cells. Values in scatter plot represent mean ± standard error of mean (s.e.m). p-values in the scatter plots are calculated with paired sample Wilcoxon signed rank test (black, one-sided) and two sample t-test (magenta and green, one-sided), with *p<5e-2, **p<1e-2, ***p<1e-3.

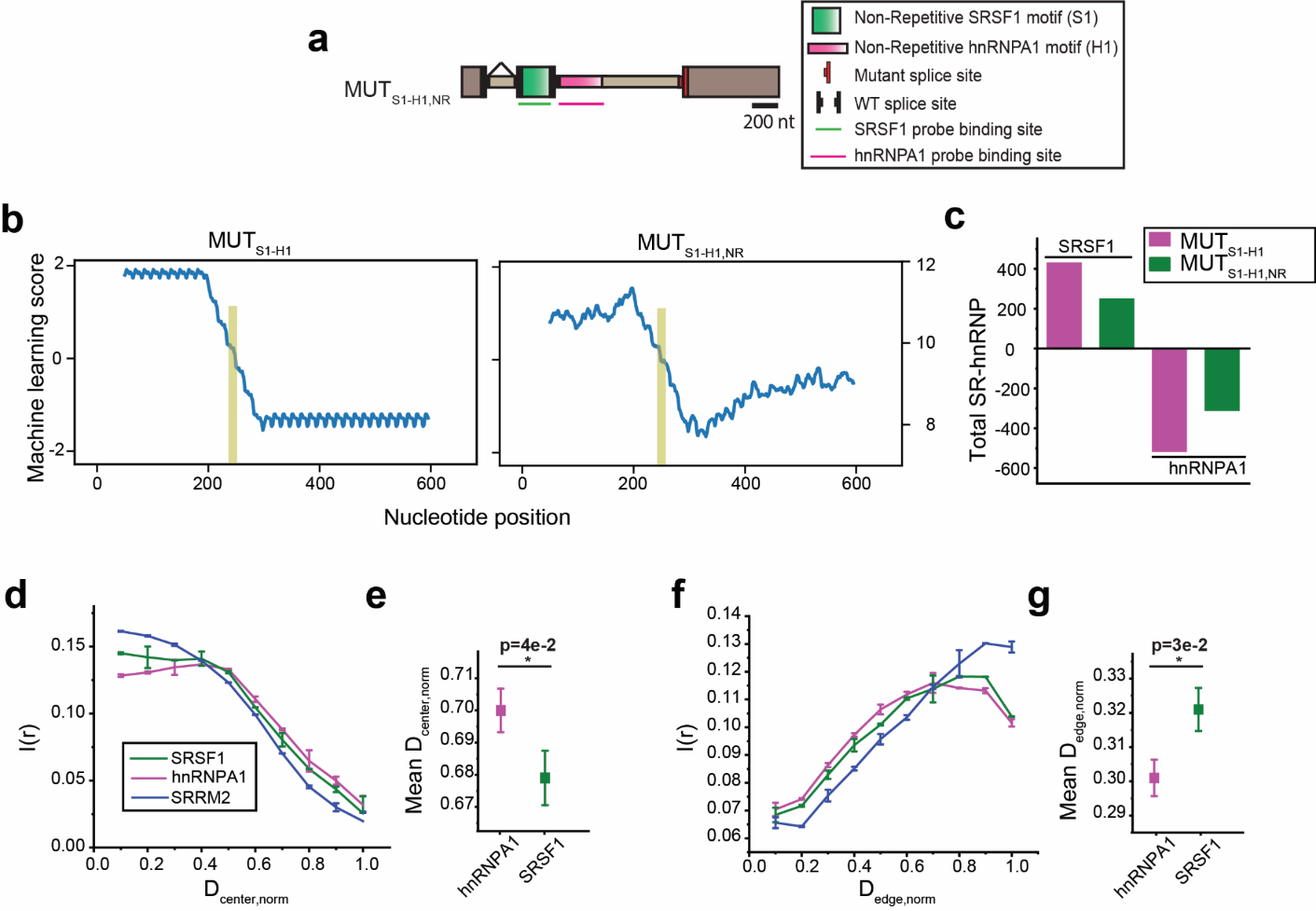

**Figure S7. Intra-speckle positioning of RNA transcripts from constructs devoid of repeats.** (a) Schematic illustration of MUT_S1-H1,NR_ construct. These constructs were obtained from MUT_S1-H1_ by mutating each nucleotide in the exonic and intronic regions with probability 37.5%. (b) Plots showing predicted SR enrichment (positive) or hnRNP enrichment (negative) for MUT_S1-H1_ and MUT_S1-H1,NR_ constructs. The prediction is based on a machine learning model trained on splicing data (*4*), and smoothed using a 100nt sliding window. Bars show predicted MaxEnt splice site score (*5*). (c) Machine learning-predicted total SR enrichment (positive) and hnRNP enrichment (negative) of the non-repeat constructs versus the original constructs. Population distribution of SRSF1 and hnRNPA1 motif signals as a function of the normalized distance from the center (d) and from the edge (f) of the speckle for MUT_S1-H1,NR_. Population-weighted mean normalized distance of SRSF1 and hnRNPA1 signal from the center (e) and from the edge (g) of speckle for each speckle in MUT_S1-H1,NR_. Error bars in the population vs. distance plots report the standard deviation from two replicates, each containing at least 60-75 nuclear speckles collected from 4-5 cells. Scatter plots are generated by combining all nuclear speckles (120-150) from two replicates. Values in scatter plot represent mean ± standard error of mean (s.e.m). p-values in the scatter plots are calculated with paired sample Wilcoxon signed rank test (one-sided), with *p<5e-2, **p<1e-2, ***p<1e-3.

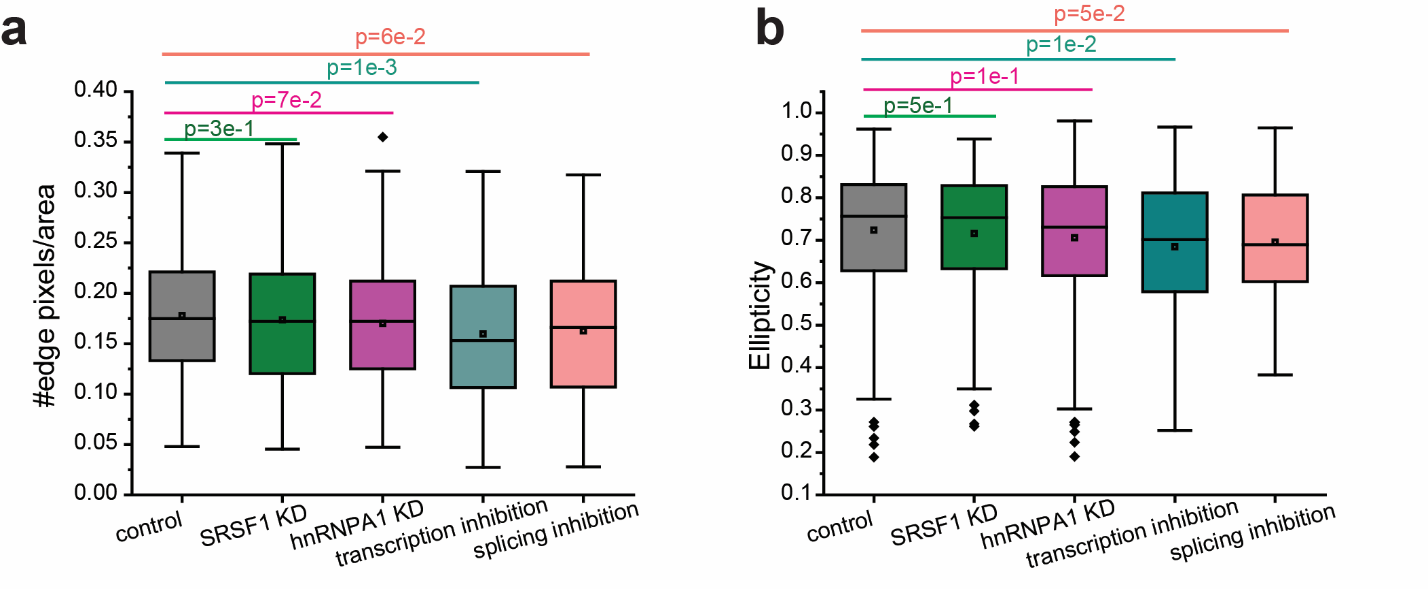

**Figure S8. The effect of siRNA mediated knockdown and drug treatments on speckle morphology.** (a) The ‘regularity’ or ‘roughness’ of the speckle surface is measured by the number of edge pixels divided by area, where number of edge pixels is an estimation of the perimeter of the speckle. The lower the value of perimeter-to-area ratio, the more regular or rounded the speckle is. (b) Ellipticity is a measure of the deviation of speckle shape from a perfect sphere and is calculated as, $\sqrt{\left( \left( a^{2}-c^{2} \right)/a^{2} \right)}$, where a is the equatorial radius and c is the polar radius assuming nuclear speckles to be an ellipse. An ellipticity value of 0 represents a perfect sphere. Each data set contains 300-400 nuclear speckles collected from 30-40 cells. p-values are calculated with two sample t-tests (two-sided). Description of box plots is same as Figure S1.

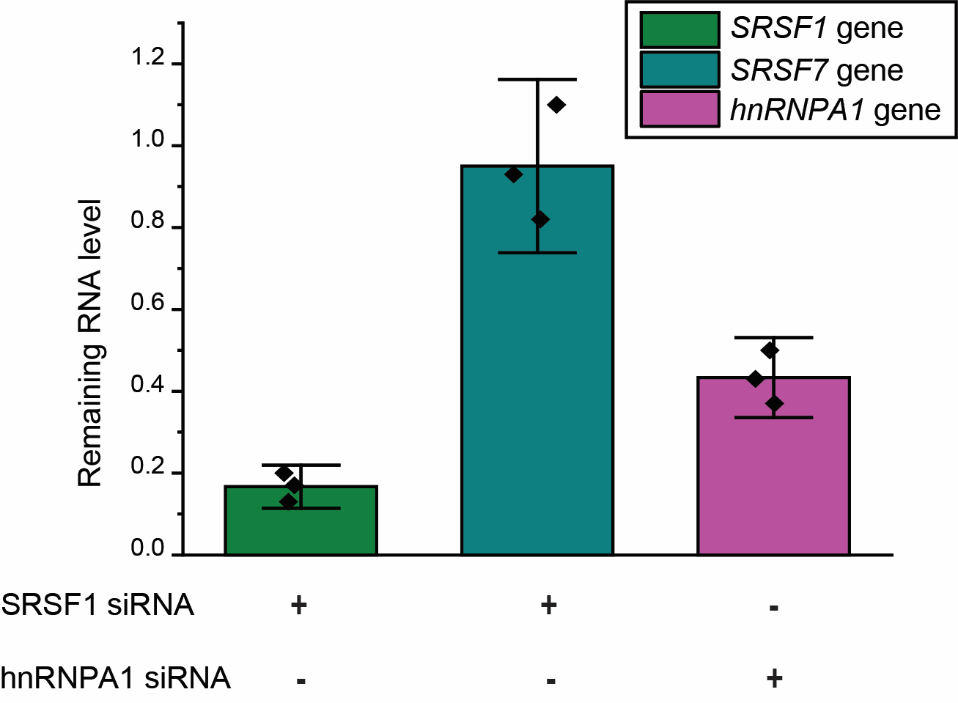

**Figure S9.** **Knockdown efficiencies at the mRNA level measured by qPCR.** The Ct value for each target gene under each knockdown condition was normalized with the Ct value of *ACTB* mRNA. The normalized Ct value of each target gene under knockdown conditions was compared to the normalized Ct value of the same gene in the control scramble siRNA. Error bars report mean ± standard deviation from three biological replicates (black points).

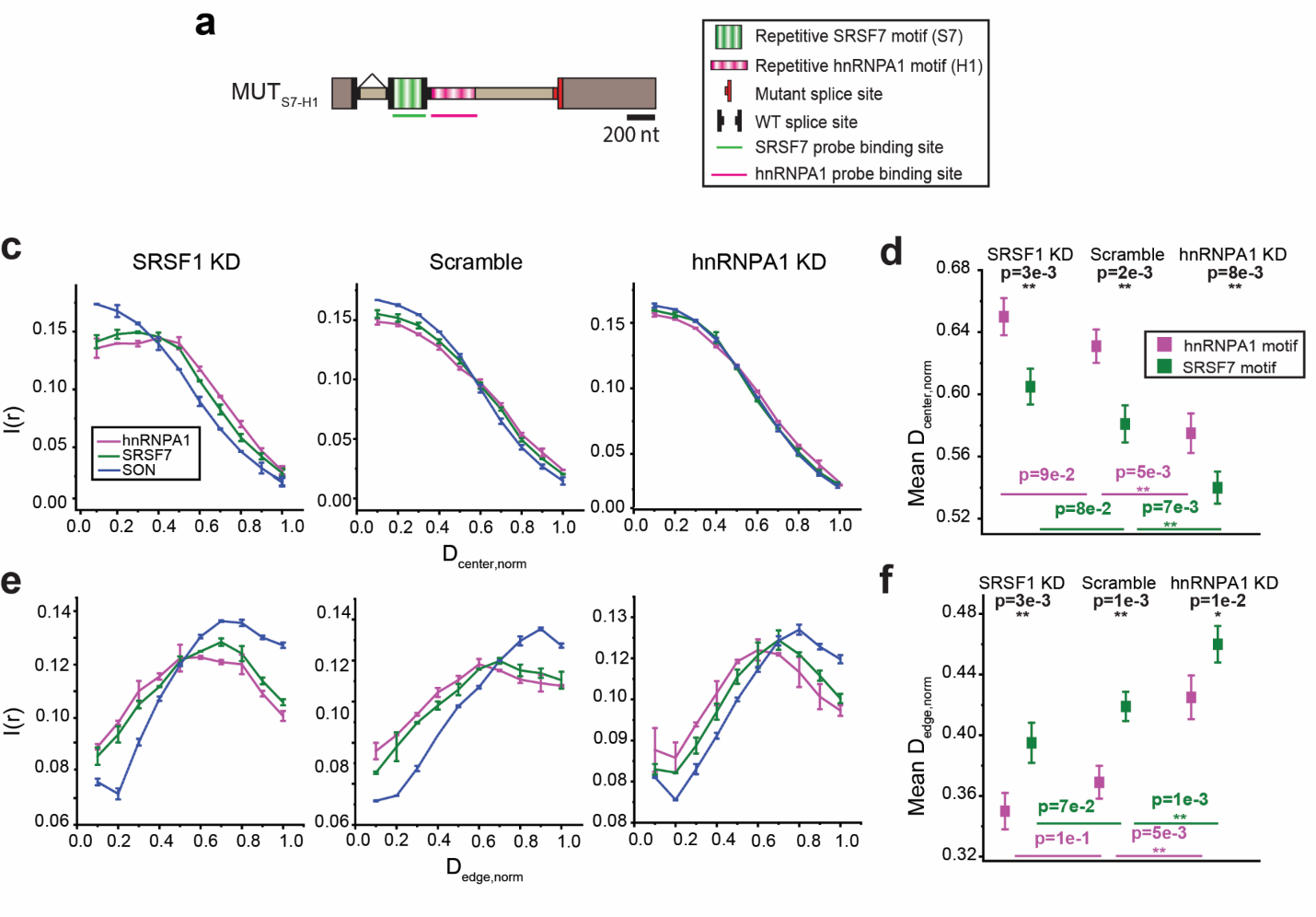

**Figure S10.** **Effect of SRSF1 and hnRNPA1 knockdown on the intra-speckle organization of RNAs containing SRSF7 motifs in exon and hnRNPA1 motifs in intron.** (a) Schematic illustration of MUT_S7-H1_. Population distribution of SRSF7 and hnRNPA1 motif signals as a function of the normalized distance from the center (b) and from the edge (d) of the speckle. Population-weighted mean normalized distance of SRSF7 and hnRNPA1 signal from the center (c) and from the edge (e) of the speckle for each speckle. Error bars in the population vs. distance plots report the standard deviation from two replicates, each containing 60-90 nuclear speckles collected from 4-6 cells. Scatter plots are generated by combining all nuclear speckles (120-180) from two replicates. Values in scatter plot represent mean ± standard error of mean (s.e.m). p-values in the scatter plots are calculated with paired sample Wilcoxon signed rank test (black, one-sided) and two sample t-test (magenta and green, one-sided), with *p<5e-2, **p<1e-2, ***p<1e-3.

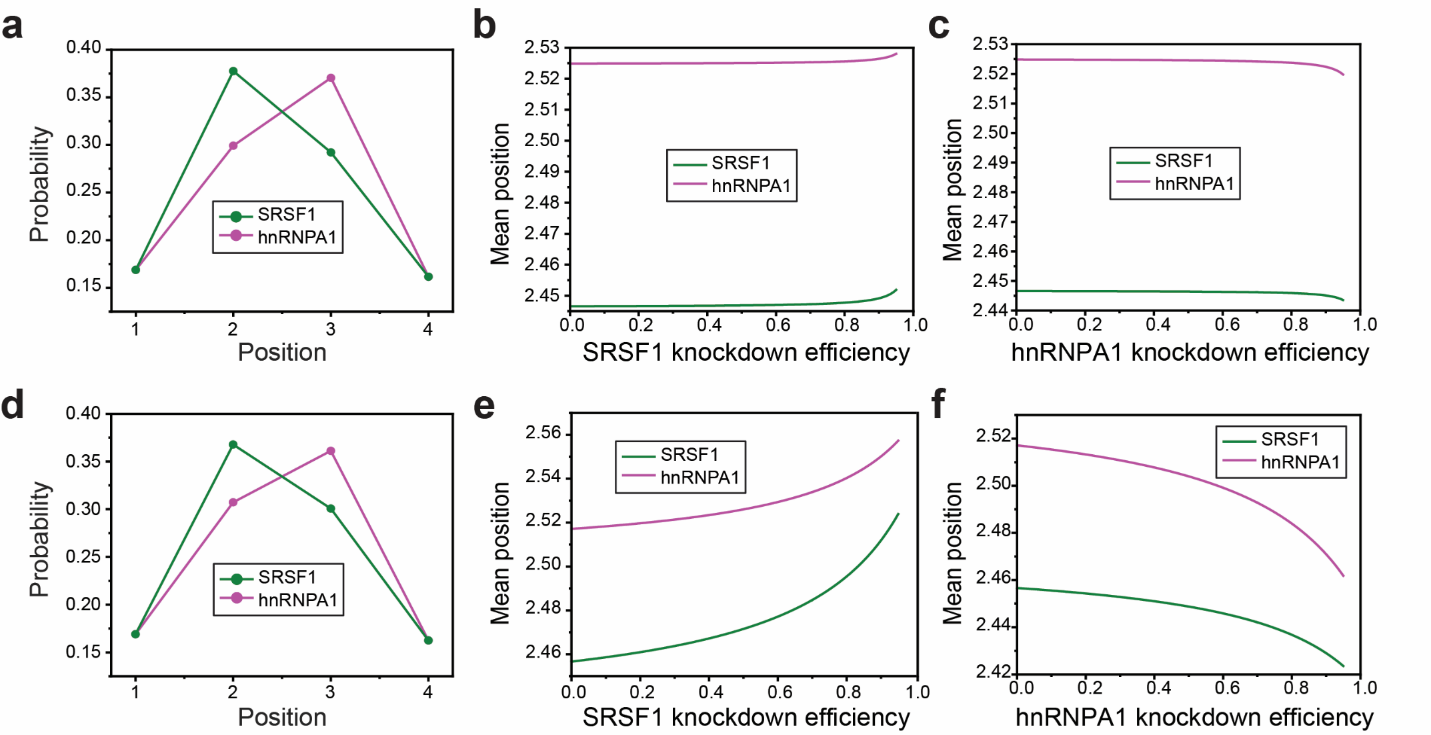

**Figure S11. Toy model for different K_d_ values. (a)-(c) K_d_ = 100 nM.** (a) Probability distributions of SRSF1 and hnRNPA1 motif position predicted by this model. Mean positions of SRSF1 and hnRNPA1 motif plotted as a function of both SRSF1 (b) and hnRNPA1 (c) knockdown efficiencies. (d)-(f) K_d_ = 10 uM. (d) Probability distributions of SRSF1 and hnRNPA1 motif position predicted by this model. Mean positions of SRSF1 and hnRNPA1 motif plotted as a function of both SRSF1 (e) and hnRNPA1 (f) knockdown efficiencies.

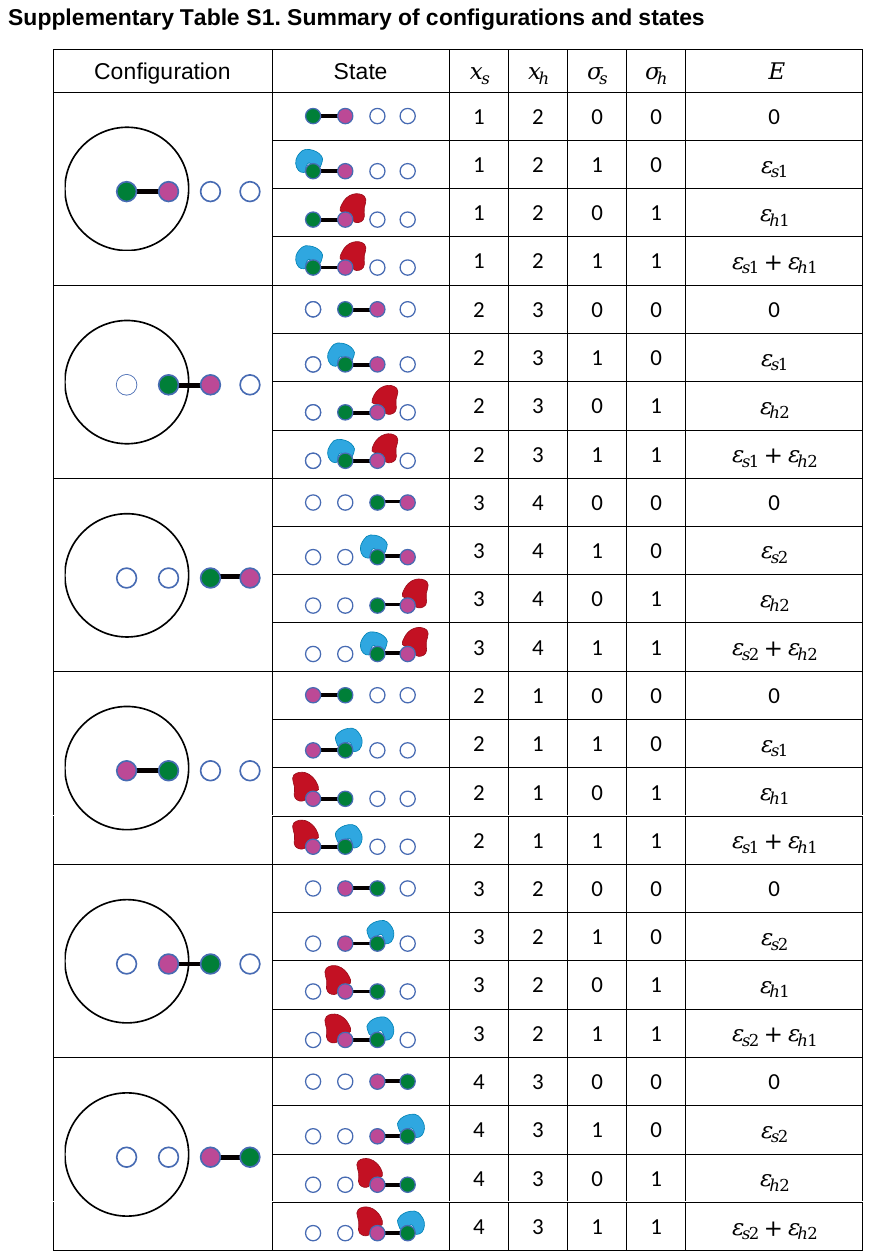

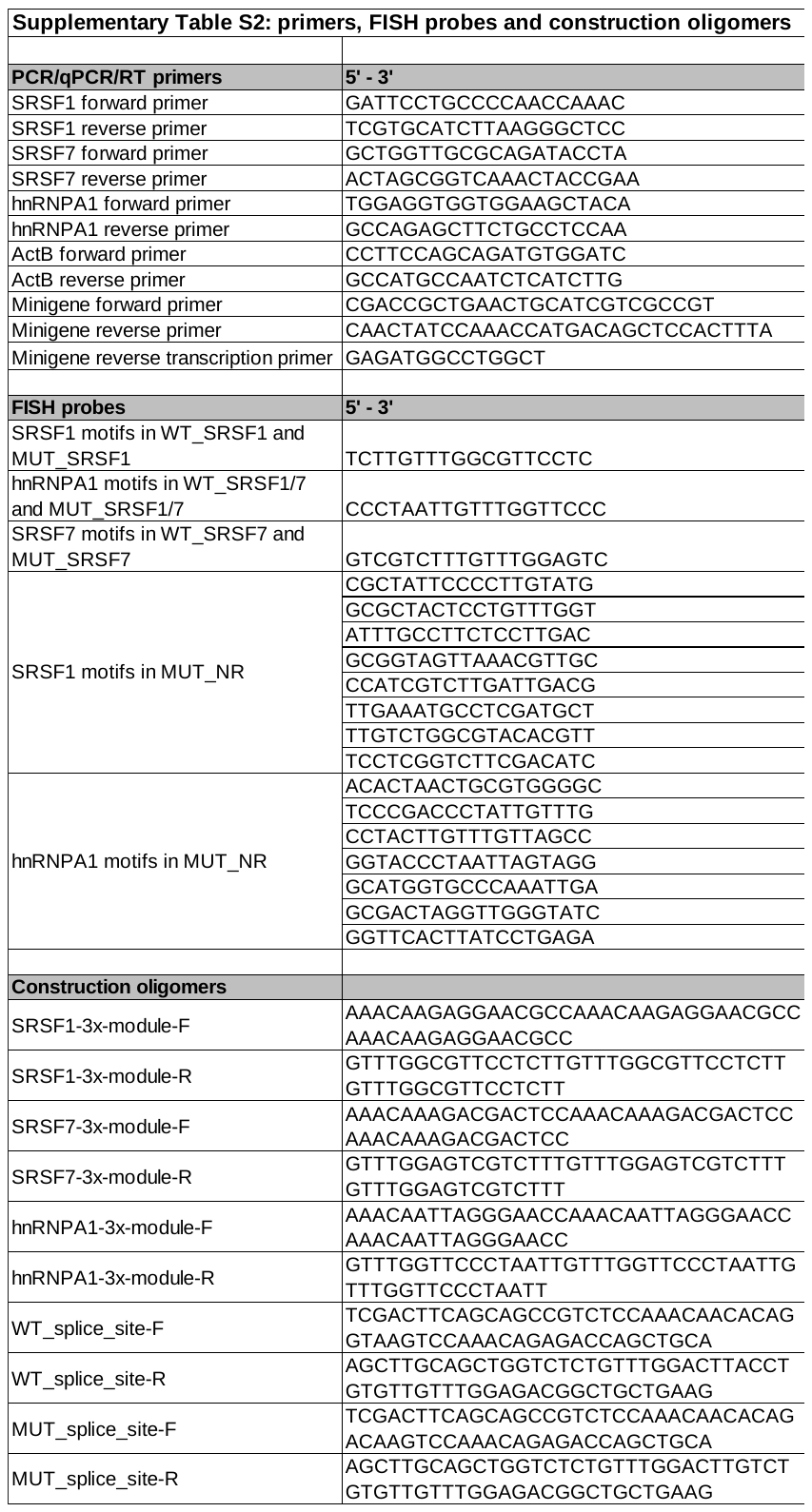

**Supplementary References**

1.	M. Y. Hein, N. C. Hubner, I. Poser, J. Cox, N. Nagaraj, Y. Toyoda, I. A. Gak, I. Weisswange, J. Mansfeld, F. Buchholz, A. A. Hyman, M. Mann, A human interactome in three quantitative dimensions organized by stoichiometries and abundances. *Cell*. **163**, 712–723 (2015).

2.	A. Fujioka, K. Terai, R. E. Itoh, K. Aoki, T. Nakamura, S. Kuroda, E. Nishida, M. Matsuda, Dynamics of the Ras/ERK MAPK cascade as monitored by fluorescent probes. *J Biol Chem*. **281**, 8917–8926 (2006).

3.	S. Maharana, J. Wang, D. K. Papadopoulos, D. Richter, A. Pozniakovsky, I. Poser, M. Bickle, S. Rizk, J. Guillén-Boixet, T. M. Franzmann, M. Jahnel, L. Marrone, Y.-T. Chang, J. Sterneckert, P. Tomancak, A. A. Hyman, S. Alberti, RNA buffers the phase separation behavior of prion-like RNA binding proteins. *Science*. **360**, 918–921 (2018).

4.	S. E. Liao, M. Sudarshan, O. Regev, *bioRxiv*, in press, doi:10.1101/2022.10.01.510472.

5.	G. Yeo, C. B. Burge, Maximum entropy modeling of short sequence motifs with applications to RNA splicing signals. *J Comput Biol*. **11**, 377–394 (2004).
